# Spatial Frequency Maps in Human Visual Cortex: A Replication and Extension

**DOI:** 10.1101/2025.01.21.634150

**Authors:** Jiyeong Ha, William F. Broderick, Kendrick Kay, Jonathan Winawer

## Abstract

In a step toward developing a model of human primary visual cortex, a recent study introduced a model of spatial frequency tuning in V1 (Broderick et al., 2022). The model is compact, using just 9 parameters to predict BOLD response amplitude for locations across all of V1 as a function of stimulus orientation and spatial frequency. Here we replicated this analysis in a new dataset, the ‘nsdsynthetic’ supplement to the Natural Scenes Dataset (Allen et al., 2022), to assess generalization of model parameters. Furthermore, we extended the analyses to extrastriate maps V2 and V3. For each retinotopic map in the 8 NSD subjects, we fit the 9-parameter model. Despite many experimental differences between NSD and the original study, including stimulus size, experimental design, and MR field strength, there was good agreement in most model parameters. The dependence of preferred spatial frequency on eccentricity in V1 was similar between NSD and Broderick et al. Moreover, the effect of absolute stimulus orientation on spatial frequency maps was similar: higher preferred spatial frequency for horizontal and cardinal orientations compared to vertical and oblique orientations in both studies. The extension to extrastriate maps revealed that the biggest change in tuning between maps was in bandwidth: the bandwidth in spatial frequency tuning increased by 70% from V1 to V2 and 100% from V1 to V3. This implies that higher visual areas are sensitive to a greater range of stimulus spatial frequencies, paralleling the fact that higher visual areas are sensitive to a greater range in location (i.e., larger receptive fields). Together, the results show robust reproducibility and bring us closer to a systematic characterization of spatial encoding in the human visual system.

## 1. Introduction

Many components of the visual system are selective to spatial scale, particularly in cortex. In both primates and cats, most V1 neurons exhibit relatively narrow bandpass tuning to spatial frequency (about 1 to 2 octaves), in contrast to the broader, low-pass tuning of LGN neurons (Campbell et al., 1969; De Valois et al., 1982; Foster et al., 1985). Preferred spatial frequency varies across neurons in V1, even within a small patch, indicating a multiscale visual representation (Campbell *&* Robson, 1968; De Valois et al., 1982). The distribution of preferred frequencies also varies across locations: for example, when averaged across local populations, peak spatial frequency decreases with eccentricity (De Valois et al., 1982; Issa et al., 2000; Movshon et al., 1978; Tootell et al., 1988; Xu et al., 2007). Spatial frequency tuning, like receptive field size, may also vary across the visual hierarchy (Vanni et al., 2020).

Electrophysiology studies provide detailed tuning curves for individual neurons and, in some cases, statistical descriptions of neural tuning within a local patch of neurons, but cannot quantify tuning across a whole map due to limited sampling. Functional MRI (fMRI) provides complementary information about spatial frequency tuning: it lacks precision at the level of individual neurons, but can systematically measure population averages across an entire map, including V1 (Broderick et al., 2022; D’Souza et al., 2016; Kay, 2011; Sasaki et al., 2001) and several extrastriate maps (Aghajari et al., 2020; Farivar et al., 2017; Henriksson et al., 2008).

The fMRI studies above all reported that preferred spatial frequency of V1 decreases with eccentricity, but differed in their absolute estimates. Notably, two recent studies, Aghajari et al. (2020) and Broderick et al. (2022) showed good agreement in estimates of peak spatial frequency, despite employing different approaches: Aghajari et al. (2020) used a voxel-wise approach, fitting independent spatial frequency tuning curves to each voxel, whereas Broderick et al. (2022) used a map-wise approach, fitting a low-dimensional model of spatial frequency and orientation tuning across an entire visual area. This agreement suggests a potential for establishing a reliable model of spatial frequency maps in human visual cortex.

Motivated by this, we sought to replicate Broderick et al. (2022). The study used “scaled grating” stimuli, in which the local spatial frequency was inversely proportional to eccentricity. These stimuli are efficient for mapping spatial frequency preferences in parallel across the visual field, because peripheral map locations are relatively insensitive to high frequencies and foveal locations are less sensitive to low frequencies. Hence the stimuli can more efficiently sample the sensitive range of each location compared to uniform gratings. A similar set of grating stimuli was used in the ‘nsdsynthetic’ supplement to the Natural Scenes Dataset (Allen et al., 2022; Gifford et al., 2026). We analyzed these data to ask how closely the findings of Broderick et al. (2022) replicated. In doing so, we adopted the same model parameterization used by Broderick et al., fitting the model to all the data in V1 simultaneously with 9 parameters. This parameterization imposes smooth variation in the model’s spatial frequency tuning as a function of voxel eccentricity, voxel polar angle, local stimulus orientation, and local stimulus spatial frequency. We evaluated the reliability of spatial frequency maps in V1 by comparing the parameters fit in the new dataset with the parameters fit to Broderick et al.’s data. Furthermore, we fit the same parameterized model to data from V2 and V3 in the new dataset.

By empirically assessing the reproducibility of Broderick et al.’s model in V1, and then applying the model to V2 and V3, we enhance our understanding of the organization of spatial frequency preferences within human retinotopic maps.

## 2. Materials and Methods

### 2.1. Datasets

No new human subjects data were collected for this project. We analyzed two datasets from published work. The original experiments were conducted in accordance with the Declaration of Helsinki and were approved by the institutional review boards at U Minnesota (NSD; Allen et al., 2022) and NYU (Broderick et al., 2022).

#### 2.1.1. NSD gratings dataset

The Natural Scenes Dataset (NSD; Allen et al., 2022) is a publicly available dataset that contains extensive fMRI responses collected at ultra-high field (7T) from eight subjects. In addition to the core NSD experiment which consisted of up to 40 sessions per subject of viewing images of natural scenes, the subjects also completed one ‘NSDsynthetic’ session (Gifford et al., 2026). This session measured brain responses to synthetic stimuli including the scaled gratings investigated in this study, as well as several other stimulus types not analyzed for this paper. In this paper, we refer to the measured responses to the scaled gratings as the ‘NSD gratings dataset’. The functional data were acquired at 7T using whole-brain gradient-echo EPI at 1.8-mm resolution. A general linear model (GLMsingle; Prince et al., 2022) was used to estimate single-trial beta weights. More details on the stimuli, data acquisition, and preprocessing can be found in the NSD papers (Allen et al., 2022; Gifford et al., 2026) and the NSD manual (see http://naturalscenesdataset.org). The preprocessed NSD data includes GLM results in multiple formats. The GLMresults used for this study is “fithrf_GLMdenoise_RR” prepared in the “nativesurface” format. These results consist of single-trial betas (expressed in units of percent BOLD signal change) for each vertex in each subject’s native cortical surface.

##### NSD experimental design

The ‘NSDsynthetic’ session consists of 8 scans, alternating between a fixation task (detect a luminance change of the fixation dot) and a one-back task on the images (detect two consecutive trials with the same image). Subjects did both tasks accurately. Percent correct in the fixation task was 89.6% ± 8.7% (mean±sd across 8 subjects) and d’ in the one-back was 2.5±0.6. For the purposes of this study, we combined the fMRI responses across the two tasks. In the occasional cases that there were consecutive repeats of a scaled grating stimulus, the second presentation was omitted from analysis. Each stimulus was presented for 2 s, followed by 2 s of blank (uniform screen, mean luminance). Each stimulus class (see next section) was presented once in each of the 8 scans. The stimuli were displayed within an 8.4° x 8.4° square centered on the screen against a gray background. Throughout the experiment, a circular anti-aliasing mask with a small semi-transparent fixation dot (0.1° radius) was present in the center of the screen. The anti-aliasing mask crops out the central region in which the spatial frequency is too high to render without aliasing. The size of the anti-aliasing mask is spatial frequency dependent, ranging from 0.02º radius for the scaled grating with the lowest frequency to 0.5º for the scaled grating with the highest frequency. Our analyses excluded vertices if the pRF center was within the largest antialiasing mask for any stimulus.

##### NSD spatial frequency stimuli

The NSD synthetic experiment contained a total of 284 images including the 112 scaled grating stimuli used in this study. These stimuli were designed to take advantage of the known inverse relationship between preferred spatial frequency and eccentricity. Scaled gratings in NSD were constructed in the same manner as the Broderick et al. (2022), according to the equation:

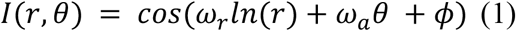

The pixel intensities in the image, *I*, at polar coordinates (*r,θ*) are specified in terms of two frequency parameters, *ω*_*a*_ (angular frequency in cycles per revolution around the image) and *ω*_*r*_ (radial frequency specifying radians per unit increase in *ln*(*r*)), and one phase parameter *ϕ* (in radians). The local spatial frequency *ω*_*l*_ depends only on the pixel eccentricity, *r*, and the frequency vector (*ω*_*r*_, *ω*_*a*_) :

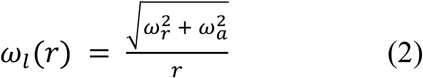

(See also Supplement section 1.1 in Broderick et al.(2022))

Likewise, the local orientation *θ*_*l*_ depends only on the pixel polar angle, *θ*, and the frequency vector (*ω*_*r*_, *ω*_*a*_):

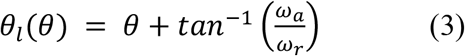

Each stimulus class was defined by its frequency vector (*ω*_*r*_, *ω*_*a*_). There were 28 stimulus classes in the experiments, with 4 exemplars per class defined by their phase (0, 1.57, 3.14, 4.71 radians). The 28 stimulus classes are organized into four shapes and “mixtures”, which are intermediate between the other four shapes (**Figure 1a**):

**Figure 1.**
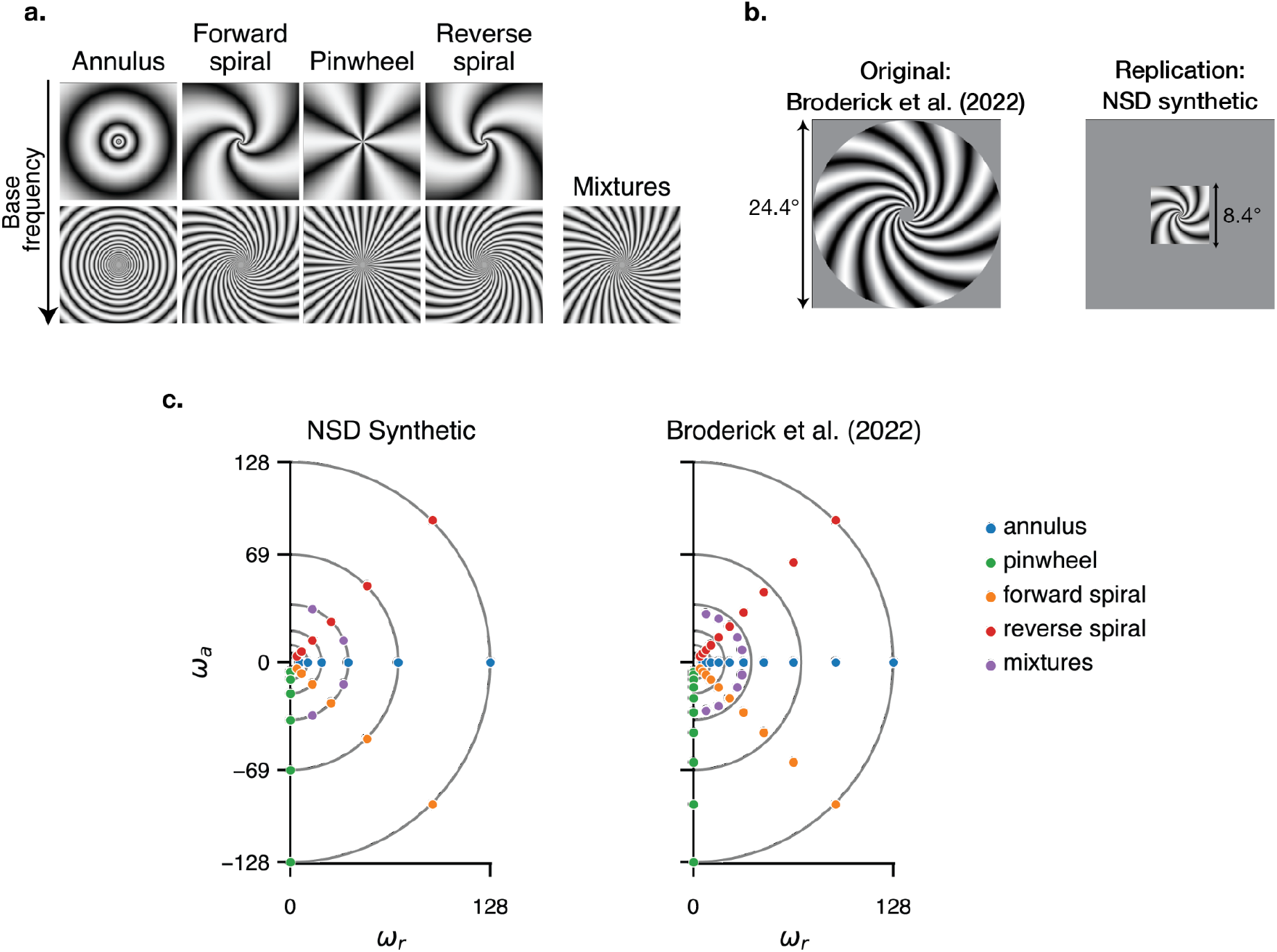
Stimuli for the two studies. (a) Example stimuli with two base frequencies (6 and 37) and five orientations (annulus, forward spiral, pinwheel, reverse spiral, mixture). Note that the mixtures were only presented at a single base frequency in each experiment. (b) Stimulus size and aperture (circular vs square) differences between the two studies. Other differences are listed in Table 1 and Supplementary Tables 2 and 3. (c) Frequency vectors (*ω*_*r*_, *ω*_*a*_) of the stimulus classes in the two studies. The gridlines are matched to facilitate comparison

1. Pinwheels: *ω*_*r*_ = 0, *ω*_*a*_ ∈ {-6, -11, -20, -37, -69, -128}
2. Annuli: *ω*_*a*_ = 0, *ω*_*r*_ ∈ {6, 11, 20, 37, 69, 128}
3. Forward spirals: *ω*_*r*_ = −*ω*_*a*_ ∈ {4, 7, 14, 26, 49, 91}
4. Reverse spirals: *ω*_*r*_ = *ω*_*a*_ ∈ {4, 7, 14, 26, 49, 91}
5. Mixtures: (*ω*_*r*_, *ω*_*a*_) ∈ {(14,-34), (34,-14), (34,14), (14,34)}

The frequency vectors were selected so that the base frequencies (vector length of (*ω*_*r*_, *ω*_*a*_)) matched across shapes (i.e., the stimulus coordinates lie on circles centred at the origin in **Figure 1c**). A constraint on the parameters is that *ω*_*a*_ are integers, as non-integer circular frequencies result in edge artifacts. Because of this constraint, there are slight discrepancies in base frequencies between the pinwheel/annulus stimuli and the forward/reverse spiral stimuli, as in Broderick et al. (2022). The frequency parameters used for each stimulus class and the resulting base and local spatial frequencies are listed in Supplementary Table 1.

##### NSD retinotopy data and visual areas

Retinotopy data were acquired from NSD subjects in a separate session. Full details are available in the original paper (Allen et al., 2022). The mapping stimuli were slowly moving apertures filled with dynamic colorful textures, extending up to 4.2 deg eccentricity. A small fixation dot (0.1 deg in radius) was superimposed on the stimuli at the screen center. A total of six scans (three with bar apertures, three with wedge/ring apertures) were conducted. Population receptive field (pRF) models were fit to the time series of each voxel from each subject. Retinotopic ROIs were hand-drawn by the NSD authors based on the analyzed pRF data. Here, we used the pRF solutions and ROIs on the individual subject native surface, as made available by the NSD.

For both datasets, we use the voxel-wise retinotopic model solutions to estimate the pRF coordinates of each voxel rather than standardized templates such as those from Benson et al. (2012; 2014), as the atlases are not yet as accurate as manually-drawn boundaries (Benson et al., 2022). Although there are many free parameters fit in the retinotopy models, these parameters are fit separately from the spatial frequency model and hence are inputs to, rather than free parameters of, the low-dimensional spatial frequency models.

#### 2.1.2. Broderick et al. dataset

We compared the results from NSD gratings dataset to those from Broderick et al. (2022), which measured BOLD responses to the scaled gratings from 12 subjects. Briefly, functional data were acquired at a 3T scanner with a spatial resolution of 2 mm. The response amplitudes to the stimuli were estimated for each surface vertex for each stimulus class, with 100 bootstraps across runs using GLMdenoise (Kay et al. 2013). The GLMdenoise algorithm differs slightly from the GLMsingle tool used for the NSD in that it estimated beta weights per stimulus type, not per trial, and in several other minor ways. We re-analyzed the data from Broderick et al. to ensure that we used the same computational methods for the two datasets. The re-analysis used the median beta weight per vertex per condition (median values across bootstrapped scans). These were made publicly available by the authors (https://archive.nyu.edu/handle/2451/63344). More detailed description of the Broderick dataset is provided in the original study.

##### Broderick et al. experimental design

Subjects participated in 12 scans, performing a one-back task on an alternating stream of black and white digits at fixation, while viewing the scaled gratings. Stimuli were constrained to a circular aperture up to 12° radius, with an antialiasing mask at the fixation center (0.96° radius). Each trial consisted of eight images with different phases from the same stimulus class, 300 ms per image with 200 ms ISI, for a total of 4 s per trial. The stimulus classes were presented in pseudo-randomized order. Subjects did the task accurately, with an average d-prime of 3.1.

#### 2.1.3. Comparison of experimental design

The scaled grating stimuli used in both NSD gratings and Broderick et al. datasets were generated in the same manner. The stimulus sets were constructed such that the maximum and minimum base frequencies were the same (**Figure 1c**). However, there were several other differences in the experimental design (**Table 1**), MR acquisition (Supplementary Table 2), and analysis pipeline (Supplementary Table 3).

**Table 1.** Differences in experimental implementation between the two datasets. The table lists the major differences in experimental design between the two studies. See Supplementary Tables 2 and 3 for differences in MR acquisition and MR analysis, respectively.

|  | Broderick et al.<br>(2022) dataset | NSD gratings<br>dataset |
| --- | --- | --- |
| Number of subjects | 12 | 8 |
| Stimulus diameter (deg) | 24.4 (circular) | 8.4 (square) |
| Number of base frequencies | 10 | 6 |
| Number of orientations | 4 | 4 |
| Number of trials per condition | 12 | 8 |
| Number of images per trial | 8 | 1 |
| Task | fixation-related | fixation-related or<br>image-related |

### 2.2. One-dimensional spatial frequency model

We fit a log-normal tuning function to the mean response amplitudes of data binned by eccentricity. For Broderick et al., the bin edges were every 1º from 1º to 11º. For NSD, the bin edges were every 0.5º from 0.5º to 4º. The different binning is due to differences in voxel resolution and stimulus field of view, both smaller in NSD. We excluded vertices whose mean response amplitude across all stimuli was negative. We express spatial frequency tuning in terms of spatial period (deg per cycle, the reciprocal of spatial frequency) rather than frequency, because the period is expected to grow linearly with eccentricity, and it is easier to assess by visual inspection how well data conform to a line than to how well data conform to an inverse linear function. The log Gaussian tuning function for eccentricity bin *b* is expressed as:

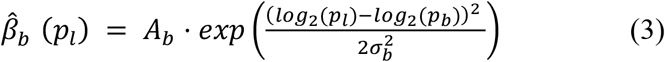

where *A*_*b*_ is the response gain, *p*_*b*_ is the preferred spatial period, *p*_*l*_ is the local spatial period (at eccentricity *b*) and *σ*_*b*_ is the bandwidth in octaves. The tuning curve was fit separately for each subject to data averaged across vertices and across the four primary stimulus shapes (annuli, pinwheels, forward spirals, and reverse spirals) for each base frequency. Hence for this model, the response of a vertex depends only on the local spatial frequency, not orientation.

The 1D model has 20 parameters per subject for Broderick et al. (bandwidth and preferred period for each of the 10 eccentricity bins) and 14 per subject per visual area for NSD, which has 7 eccentricity bins. We summarized these 1D model parameters across subjects using a precision-weighted average. Each subject’s contribution to the average was weighted by the reciprocal of the variance,

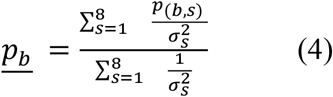

where *p*_(*b,s*)_ is the peak spatial period for eccentricity bin *b* and subject *s*, and 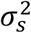 is the variance for subject *s*, defined as follows:

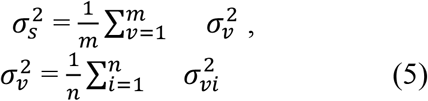

The subject variance, 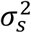 is the variance averaged across the *m* vertices in a map. The variance of each vertex, 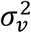, is the variance averaged across 24 stimulus classes, with the variance for the *i*^th^ stimulus class, 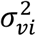, computed across beta weights for the 8 repeated images within each stimulus class *i*. The precision-weighted average assigns greater weight to data with high signal-to-noise ratio, while still incorporating all of the data. Because each subject is assigned a single precision value, the weight of each subject is the same for every eccentricity bin.

### 2.3. Two-dimensional spatial frequency model

Since the one-dimensional tuning curve model cannot measure the effect of stimulus orientation and retinotopic angle, Broderick et al. (2022) further developed a two-dimensional model for individual vertex responses. The model takes inputs as the vertex eccentricity and polar angle, and the stimulus spatial frequency and orientation at that location, and predicts the BOLD response as output (**Figure 2**). We can thus express the model response to a given stimulus for vertex *v* as follows:

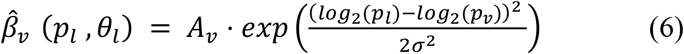

**Figure 2.**
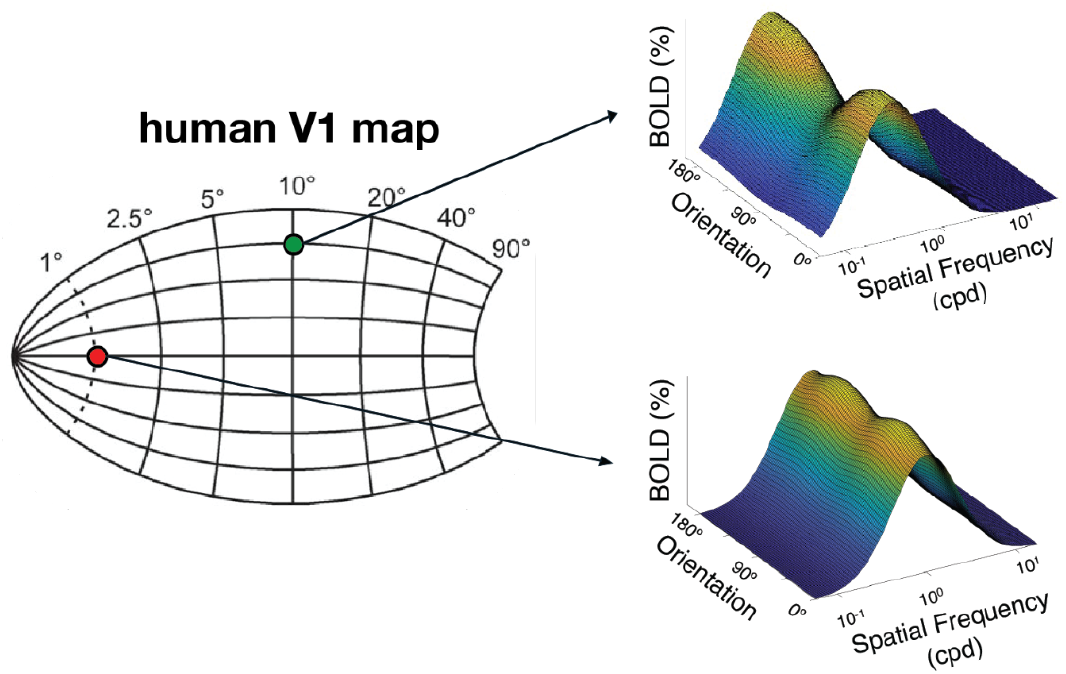
2D model schematic. Each location in V1 (left) corresponds to a visual field position, specified by eccentricity and polar angle. At each location, the model (right) has a 2D tuning curve, predicting the BOLD amplitude as a function of local spatial frequency and orientation. The schematic shows example tuning curves for two locations in the visual field (red dot: 1º eccentricity on the horizontal meridian; green dot: 10º eccentricity near vertical meridian). The peak spatial frequency is higher for the more foveal location and the orientation modulation is larger for the other location.

Here *p*_*v*_ is preferred period of the vertex and *σ* is the bandwidth, assumed to be equal across the map. *p*_*l*_ is the local spatial period (degree/cycle) and *θ*_*l*_ is the local orientation (radians). To understand how spatial frequency preferences vary across vertices within a visual area, *p*_*v*_ is forced to linearly vary as a function of vertex eccentricity *r* _*v*_: *p*_*v*_ = *m* ⋅ *p*_*v*_ + *b* . Furthermore, *p*_*v*_ is modulated by orientation. Specifically, *p*_*v*_ is expressed as:

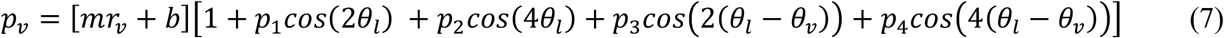

Each of the parameters *p*_*i*_ controls how *p*_*v*_ depends on a particular aspect of orientation. Specifically, for positive values of *p*_*i*_, preferred period is larger for some orientations than others:

1. *p*_1_: Vertical > horizontal (“horizontal effect”)
2. *p*_2_: Cardinal > oblique (“oblique effect”)
3. *p*_3_: Annulus > pinwheel (“radial effect”)
4. *p*_4_: Non-spirals (annuli/pinwheels) > spirals (“spiral effect”)

Those four *p*_*i*_ can be grouped into absolute orientation effects (*p*_1_ and *p*_2_) and relative orientation effects (*p*_3_ and *p*_4_). Absolute orientation effects depend on the local orientation *θ*_*l*_ but not on the vertex’s retinotopic angle *θ*_*v*_. Relative orientation effects depend on the difference between the local orientation *θ*_*l*_ and a vertex’s retinotopic angle *θ*_*v*_.

Similarly, we allow the amplitude of the BOLD response to vary with stimulus orientation:

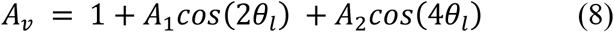

Following Broderick et al. (2022), we do not allow amplitude to vary with relative orientation, as their cross-validation analyses showed that including these parameters did not improve model accuracy. Their 9-parameter model provided the best fit among several alternatives with fewer or more parameters. Here, we use the same 9-parameter model in order to compare parameter estimates between datasets. We summarize the 9 model parameters in **Figure 3**.

**Figure 3.**
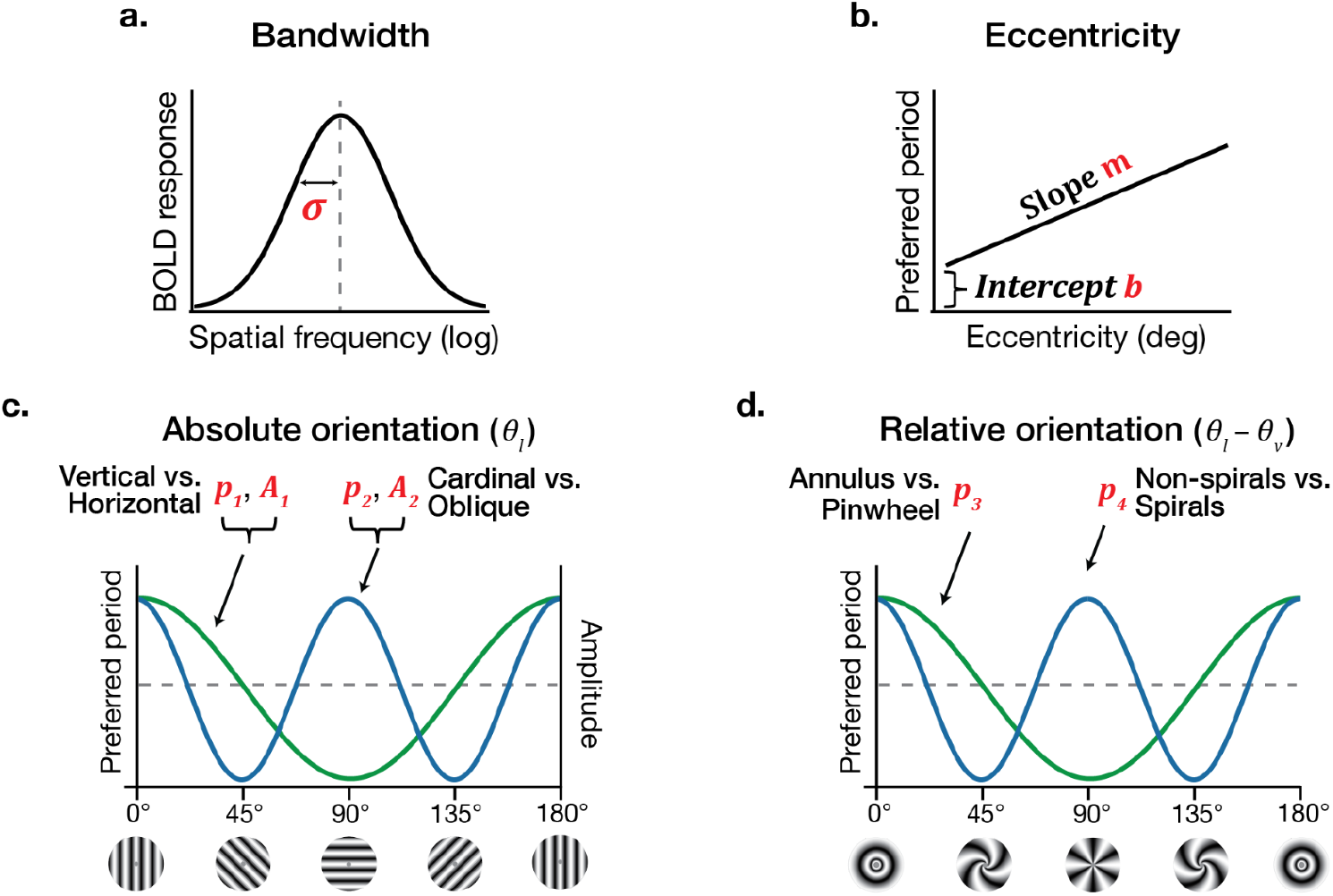
Schematic representation of the 9 parameters in the two-dimensional model. (a) Bandwidth *σ*. In this study, *σ* is assumed to be consistent across eccentricity, retinotopic location, and local stimulus properties. (b) Eccentricity effect (Slope *m* and intercept *b*). Preferred period is assumed to be an affine function of eccentricity. (c) Four parameters related to absolute orientation. *p*_3_ and *p*_4_ represent the orientation effect on preferred period, whereas *A*_3_ and *A*_4_ represent the effect on the gain of the BOLD responses. *p*_3_ and *A*_3_ are positive when the presented orientation is vertical vs. horizontal. *p*_4_ and *A*_4_ are positive when the orientation is cardinal vs. oblique. (d) Two parameters on the stimulus orientation relative to retinotopic angle. *p*_3_ is positive when the relative orientation is annulus (radial) vs. pinwheel (tangential) and *p*_2_ is positive when the orientation is spirals vs. non-spirals.

We assessed model accuracy as the cross-validated loss across several models (**Supplementary Figure 1a**). Unlike Broderick et al., in this dataset, the 9-parameter model does not have the smallest loss. However, parameter estimates were very stable across all models tested other than two highly reduced models which excluded either slope or intercept of preferred period as a function of eccentricity (**Supplementary Figure 1b**). Hence the parameter estimates we derived from fitting the 9-parameter model are unlikely to be artifacts of overfitting a model with too many free parameters.

We fit the two-dimensional model to all data within an entire map at once, excluding vertices whose pRF centers were more than one pRF size outside a 4.2º radius, and vertices whose averaged responses across the 28 stimulus classes were negative.

Following Broderick et al., the loss function was weighted by vertex precision, giving less weight to noisier vertices. The loss for vertex *v* is expressed as follows:

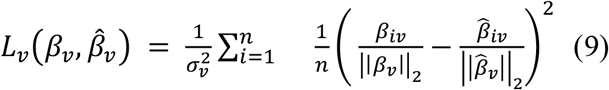

Here the loss, *L*_*v*_, is the weighted, L2-normalized mean-squared error between beta weights from the GLM and predicted responses from the model. The 28 stimulus classes are indexed by *i. β*_*iv*_ is the beta estimate of voxel *v* to stimulus class *i* from the GLM. 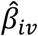 is the model prediction for stimulus class *i* and vertex *v*. ||*β*_*v*_||_2_ is the L2-norm of *β*_*v*_ across the 28 stimulus classes, and *σ* is the vertex variance, computed by taking the mean of the variance computed separately for each of the 28 stimulus classes (8 trials per class).

All model fitting procedures, including the one-dimensional tuning curves and the 2D model, were written in PyTorch (Paszke et al., 2019) using the Adam optimization algorithm (Kingma *&* Ba, 2014).

### 2.4. Bootstrapping and confidence intervals

Parameter estimates are reported as precision-weighted means across the full set of subjects (8 for NSD, 12 for Broderick et al.). Confidence intervals are derived from bootstrapping from across subjects.

For the 1-D model, tuning curves were fit separately to each eccentricity bin in each subject, as described above. Then, for each bootstrap, we selected *n* subjects at random with replacement, where *n* is the number of subjects (8 or 12), and computed the subject-wise precision-weighted mean for each parameter (bandwidth and preferred period for each eccentricity bin). This was repeated 1,000 times. The 68% confidence interval for each parameter was reported as the 16th and 84th percentile value of that parameter from the 1,000 bootstraps. The 95% confidence intervals were reported as the 2.5th and 97.5th percentile from the 1,000 bootstraps.

Effectively, the same procedure was used for the 2D model. The 9-parameter model was solved separately for each subject using all of the data from that subject (except when cross-validating, as reported in the supplement). For each bootstrap, we randomly selected *n* subjects with replacement, and computed the subject-wise precision-weighted mean for each of the 9 parameters (except for a control analysis with equal weighting across subjects reported in the supplement). This was repeated 1,000 times. The 68% and 95% confidence intervals for each parameter were derived in the same way as for the 1D model, i.e., from the appropriate percentile value for each parameter from the 1,000 bootstraps.

When comparing bandwidth among V1, V2, and V3 in the 2D model, pairwise differences were computed within each of the 1,000 bootstraps, e.g., *σ*_*V2*_ minus *σ*_*V1*_. This yields 1,000 estimates of the parameter differences from which to derive the 68th and 95th percentile confidence intervals.

### 2.5. Comparing models across datasets

To quantify the similarity of model parameters across datasets, we converted each of the nine parameters to effect sizes (Cohen’s *d*), dividing the mean estimate by its pooled standard deviation across datasets. Effect size is meaningful only when zero is a principled baseline; this holds for all parameters except bandwidth *σ*, which we reparameterized as its reciprocal (1/*σ*). The reciprocal approaches zero when there is no eccentricity tuning, providing the necessary baseline. We summarized cross-dataset agreement by the squared error between each pair of parameter estimates, and by the mean squared error (MSE) across the 9 parameter pairs. The MSE captures the replicability of the parameter profile, giving more weight to parameters that are large relative to inter-subject variability, similar to the approach in the Open Science Collaboration (2015).

We compared each of these metrics to those obtained from null distributions constructed from the NSD data using three targeted permutation procedures. Because the nine model parameters capture three distinct response properties — orientation selectivity (*p*_1_–*p*_2_, *A*_1_–*A*_2_), eccentricity dependence (slope *m*, intercept *b*), and spatial frequency tuning (bandwidth *σ*) — each procedure selectively disrupts one property while leaving the others intact, allowing us to construct a targeted null distribution for each parameter subset. We then re-solved the model for each of 1,000 permutations.

To derive null distributions for the orientation parameters, we shuffled orientation labels within each of the six base spatial frequencies and across the four mixture stimuli, preserving spatial frequency assignments; as expected, this had little effect on bandwidth, slope, or intercept, while the orientation parameters centered near zero. To null the eccentricity parameters, we shuffled eccentricity assignments across voxels while leaving orientation labels intact; bandwidth and orientation parameters were largely unchanged, the slope centered near zero, and the intercept near the mean preferred spatial frequency. To null bandwidth, we shuffled spatial frequency labels across the 28 stimulus classes; the resulting inverse bandwidth values (1/σ) were small but did not center exactly at zero, reflecting the non-negativity constraint on this parameter.

The three permutation procedures were combined into a single 9×1,000 null array — orientation parameters from the first, eccentricity parameters from the second, bandwidth from the third — against which we compared the parameters estimated from Broderick et al.

## 3. Results

The ‘nsdsynthetic’ supplement to the Natural Scenes Dataset (Allen et al., 2022; Gifford et al., 2026) included scaled grating stimuli, similar to those used by Broderick et al. (2022), but with several differences in the stimulus and experiment details (**Table 1**). Notably, due to constraints in the display setup, NSD stimulated less of the visual cortex than Broderick et al. (3x lower eccentricity extent) with fewer trials (224 vs 733). On the other hand, the NSD used smaller voxels, higher field strength, and smaller image pixels, enabling measurements closer to the fovea. Before fitting the full 2D model, we did a coarse analysis on data binned across vertices (within an eccentricity band) and binned across stimulus orientation. This analysis fits 1D spatial frequency tuning curves to data averaged within eccentricity bins.

To preview the results, we first show with the 1D model fits that NSD has good quality spatial frequency data, both in V1 and in extrastriate maps (V2 and V3). The V1 data are in reasonably good agreement with Broderick et al. We then examine the V1 data in more detail, comparing 2D model fits between NSD and Broderick et al. Most but not all parameters are in good agreement between the two datasets. Finally, we report the 2D model fits in NSD V2 and V3. The biggest difference between tuning in these areas in V1 is larger bandwidth in the extrastriate maps.

### 3.1. 1D model: NSD-synthetic has good responses to scaled grating stimuli

The 1D tuning curve analysis shows high quality data in NSD. We first consider an example dataset from one subject (**Figure 4**). For this subject, the BOLD responses in V1, V2, and V3 were all well fit by log-Gaussian tuning curves, similar to Broderick et al. As expected, the preferred spatial frequency declined with eccentricity: in all three visual maps, the tuning curves peaked at about 2 to 3 cpd in the foveal bins (0.5 to 1.0 deg), and about 1 cpd in the parafoveal bins (3.5 to 4 deg). The bandwidth was also narrower in V1 than in V2 and V3.

**Figure 4.**
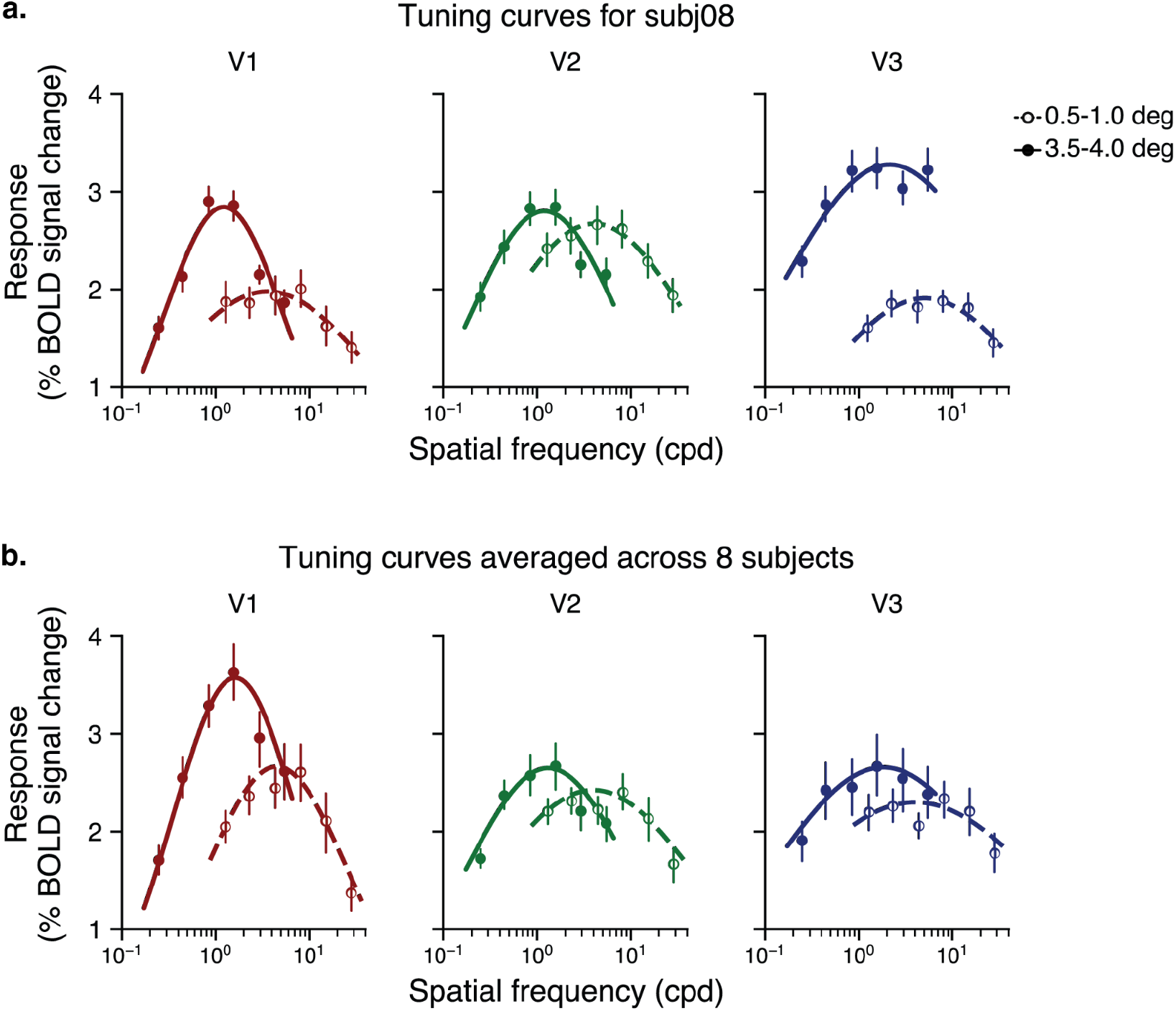
Spatial frequency tuning curves for NSD. (a) Spatial frequency tuning curves for an example NSD subject (subj08) in V1, V2, and V3. Data are plotted for two eccentricity bins. The foveal data peak at a higher spatial frequency. Error bars represent 68% confidential interval bootstrapped across trials. (b) Same as panel A but averaged across the 8 NSD subjects, with error bars representing the 68% confidence intervals bootstrapped across subjects.

Next we quantified these observations across all subjects and eccentricity bins. We found that the data show two expected systematic patterns, in general agreement with Broderick et al.

First, in V1, the preferred period (reciprocal of preferred spatial frequency), increases with eccentricity, and is well described by a straight line with a positive intercept (**Figure 5a**). The preferred period in NSD is similar to that in Broderick et al., but with a slight tradeoff in that NSD has a lower intercept and slightly higher slope. There is good agreement in preferred period between NSD observers, as evident by the small confidence intervals in **Figure 5a**.

**Figure 5.**
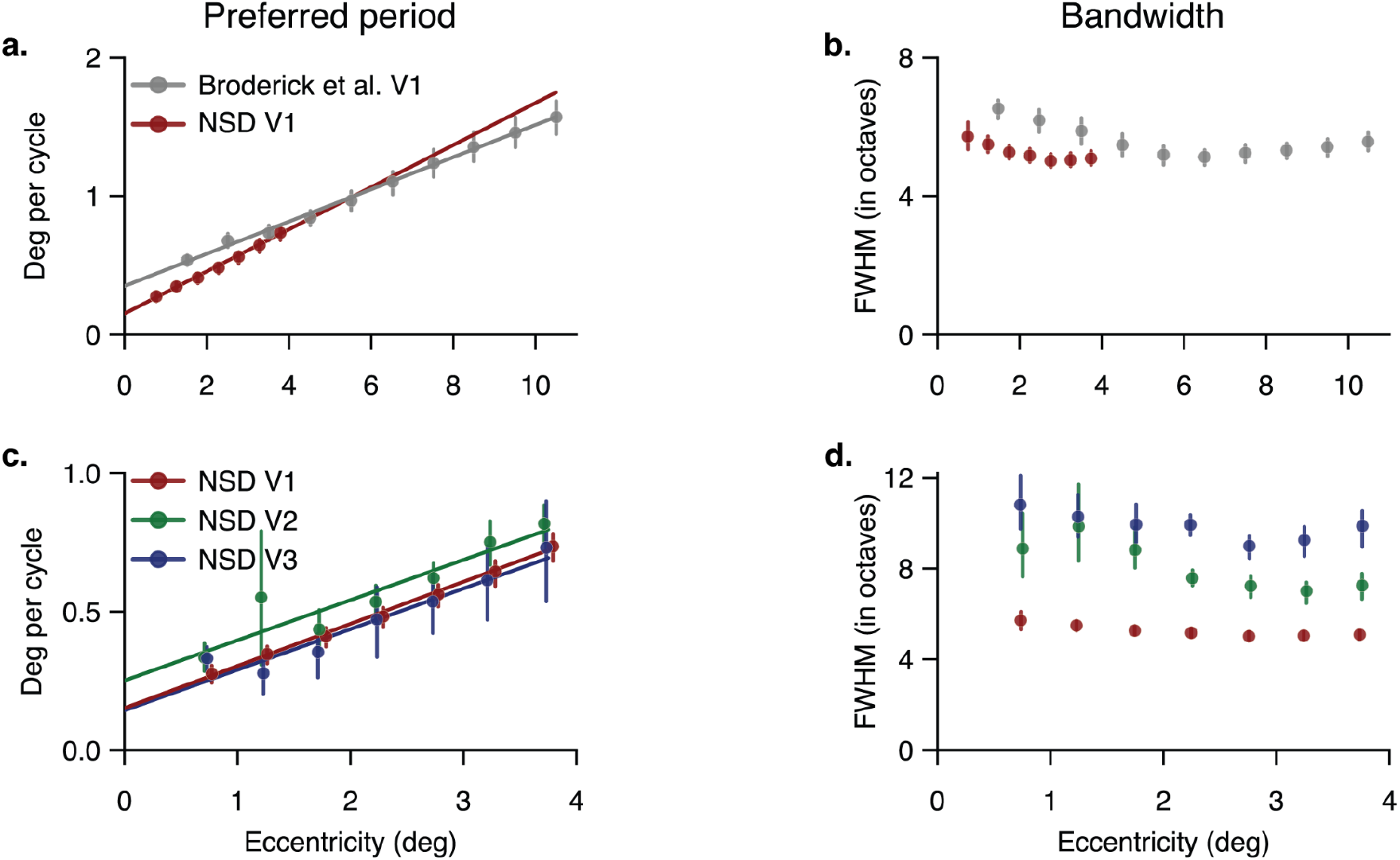
Preferred period (left column) and bandwidth (right column) for Broderick et al. and NSD, estimated by 1D model. The top row (a,b) shows V1 properties in Broderick et al. and NSD (“Replication”) and the bottom row (c,d) shows results from V1, V2 and V3 in NSD, with NSD V1 data replotted (“Extension”). (a,c) Preferred period of tuning curves as a function of eccentricity. (b,d) Full-width at half max (in octaves) of tuning curves as a function of eccentricity. Circles indicate precision-weighted means at each eccentricity bin and lines are the best linear fits to the data points. The error bars represent 68% bootstrapped confidence intervals across subjects.

Second, the bandwidth of the tuning curves in NSD V1 is large, about 5 octaves (full width at half max) for eccentricities over 2 deg, and a little higher near the fovea. This is similar to the pattern in Broderick et al’s dataset (**Figure 5b**), although the bandwidth in NSD is slightly lower.

We also examined spatial frequency tuning in V2 and V3 in NSD, which was not reported by Broderick et al. We find that V2 and V3 show reliable increases in preferred period with eccentricity (**Figure 5c**). As in V1, these data are well fit by lines with positive intercepts. The agreement across NSD subjects in V2 and V3 is not quite as high as in V1, as indicated by the larger error bars. The tuning width of NSD substantially increases from V1 to V2, and slightly more from V2 to V3 (**Figure 5d**), paralleling the tendency for receptive field size to increase along the cortical hierarchy (Dumoulin *&* Wandell, 2008). Averaged across eccentricity bins, the tuning width (full-width at half max in octaves) in NSD increases from about 5 in V1 to 8 in V2 to 10 in V3 (More precise estimates are reported for the 2D model in subsequent sections).

### 3.2. A comparison of 2D model fits for NSD V1 vs Broderick et al. V1

To compare the NSD and Broderick et al. results more quantitatively, we used the same two-dimensional model from Broderick et al. and fit it to each NSD subject’s data. The model predicts the BOLD response amplitude at each location in the V1 map as a function of the local spatial frequency and orientation in the stimulus, where “local” means the value at the pRF center. The model predictions differ across locations (i.e., vertices) but in highly constrained ways, enabling us to make reasonably good predictions for all stimulus orientations, spatial frequencies, and map locations with only 9 free parameters. Thus, the model provides a concise description of spatial frequency preferences in an entire visual field map. To ensure that any differences in parameter estimates are due to differences in the datasets rather than the analysis pipeline, we re-analyzed the Broderick et al. dataset using the same code applied to the NSD. The re-analysis produced nearly the same results.

We also report results of several simpler models which omit some of the parameters in **Supplementary Figure 1**. The parameter estimates from the simpler models are close in agreement with those from the 9-parameter model, with the exception of 2 highly reduced models.

Below, we compare the 9 parameter estimates between NSD and Broderick et al. as in Broderick et al., we combined model parameters across subjects using a precision-weighted average (see Methods). To preview the results, we find that parameters which pertain to eccentricity and to absolute orientation (stimulus orientation irrespective of pRF location) generally replicate well, whereas the two parameters that pertain to stimulus orientation relative to the pRF polar angle do not.

#### 3.2.1. V1 spatial frequency bandwidth is similar between NSD and Broderick et al

Because the 1D model showed that the bandwidth was relatively stable across eccentricity bins, the 2D model assumed a single bandwidth (in octaves, not in degrees). The bandwidth parameter *σ* was the same for NSD and Broderick et al., 2.2±0.1 and 2.3±0.1 octaves, respectively (**Figure 6 left**). This value is the standard deviation of the Gaussian tuning curve fit to log spatial frequency data. A standard deviation of 2.2 corresponds to a full-width-at-half-max of 5.2, nearly identical to the value of 5.3 estimated from the 1D model for V1 (**Figure 5b**). To visualize model tuning curves with the fit widths, we plot example model tuning curves (**Figure 6 right**).

**Figure 6.**
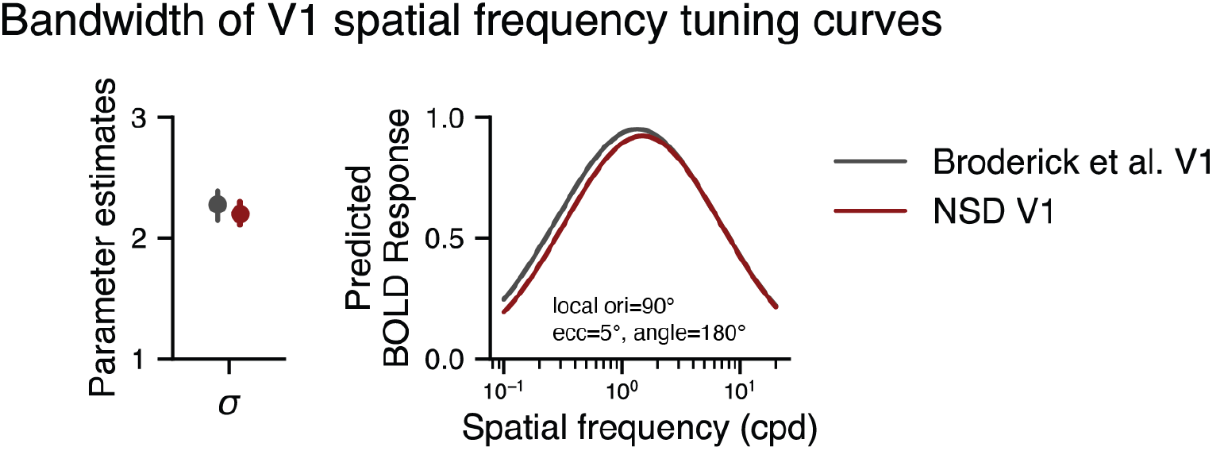
Bandwidth of V1 spatial frequency tuning curves (“Replication”). The left plot shows the estimated sigma parameter for the two datasets, Broderick et al. and NSD. The circles represent precision-weighted averages of bandwidth, where bandwidth is one standard deviation of the log Gaussian tuning curve, in octaves. The error bars indicate 68% bootstrapped confidence intervals across subjects. The right panel shows example model responses, averaged across subjects, assuming local orientation at 90° and a voxel’s pRF eccentricity and angle at 5° and 180°, respectively.

#### 3.2.2. V1 spatial frequency tuning as a function of eccentricity is similar for NSD and Broderick et al

Next we compared the effect of eccentricity on preferred period between the two datasets in V1. The slope parameter *m* showed good agreement: Broderick et al. had an average slope of 0.12±0.02, while NSD V1 had a slope of 0.15±0.01 (**Figure 7 left**). However, there was a noticeable difference in the intercepts *b*, with NSD intercept half of Broderick et al.’s (NSD 0.17±0.03 vs Broderick et al. 0.37±0.05). The lower intercept for NSD may be due to the differences in experimental details, such as the voxel size or stimulus size. Nonetheless, both intercept values were reliably above zero, indicating that spatial period is not proportional to eccentricity. Moreover, across eccentricities, the lower intercept but slightly higher slope in NSD compared to Broderick et al. trade off, such that the models’ preferred period differs by no more than 0.2 deg in the central 10 deg of eccentricity (**Figure 7 right**). These differences are small relative to the range of spatial frequency preferences reported in the literature (**Supplementary Figure 4**).

**Figure 7.**
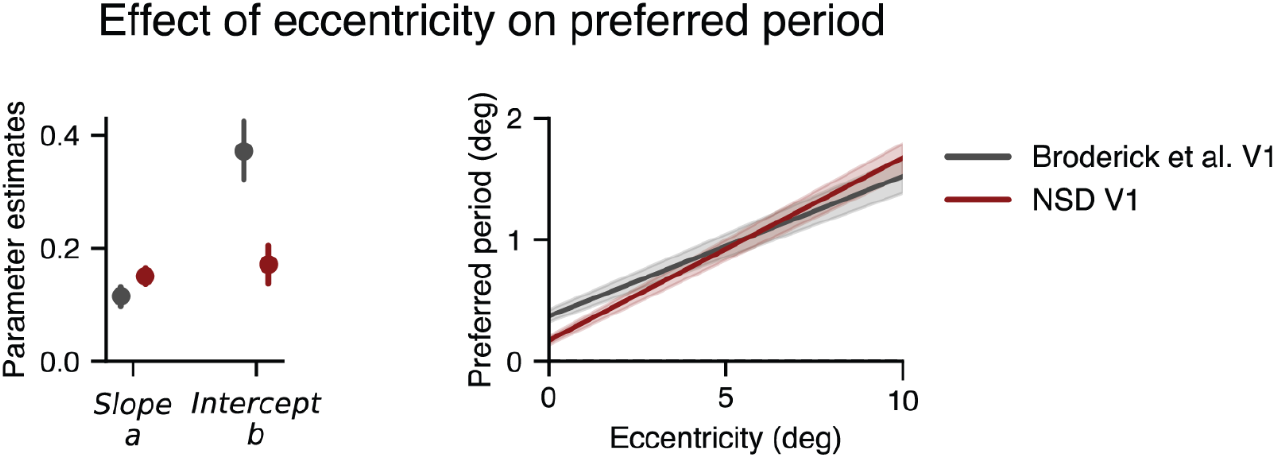
Eccentricity effect of V1 in Broderick et al. and NSD (“Replication”). The left panel shows the estimated slope and intercept from the two-dimensional model. The circles represent precision-weighted averages. The error bars indicate 68% bootstrapped confidence intervals across subjects. The right panel shows the predicted preferred period as a function of eccentricity based on the values shown left, with local orientation at 90° and a voxel’s retinotopic angle at 180°.

#### 3.2.3. Both V1 datasets show slightly higher preferred period for vertical than horizontal and for oblique than cardinal

Both datasets show that stimulus orientation modulates spatial frequency tuning. This is shown by the estimates of parameters *p*_1_ and *p*_2_. The *p*_1_ estimates are similar across the two datasets and indicate a larger preferred period for vertical than horizontal orientations (**Figure 8a**): 0.07±0.02 and 0.08±0.04 for Broderick et al. and NSD, respectively. The *p*_2_ parameters are also similar and because they are negative, indicate a larger preferred period for oblique than cardinal orientations (**Figure 8b**): -0.03±0.006 and -0.02±0.01 for Broderick et al. and NSD, respectively. The negative value of *p*_2_ aligns with the well-documented “oblique effect” in visual perception: poorer discrimination and detection for oblique than cardinal orientations (Appelle, 1972; Mach, 1861). It is not clear whether the positive value of *p*_1_ has a psychophysical correlate, as performance differences between horizontal and vertical stimuli are inconsistent across studies (for example, Berkley et al., 1975; Bulens et al., 1988; Campbell et al., 1966; Long *&* Tuck, 1991). Both of these orientation effects are modest in size, indicating a variation in preferred period as a function of orientation of no more than about 12%.

**Figure 8.**
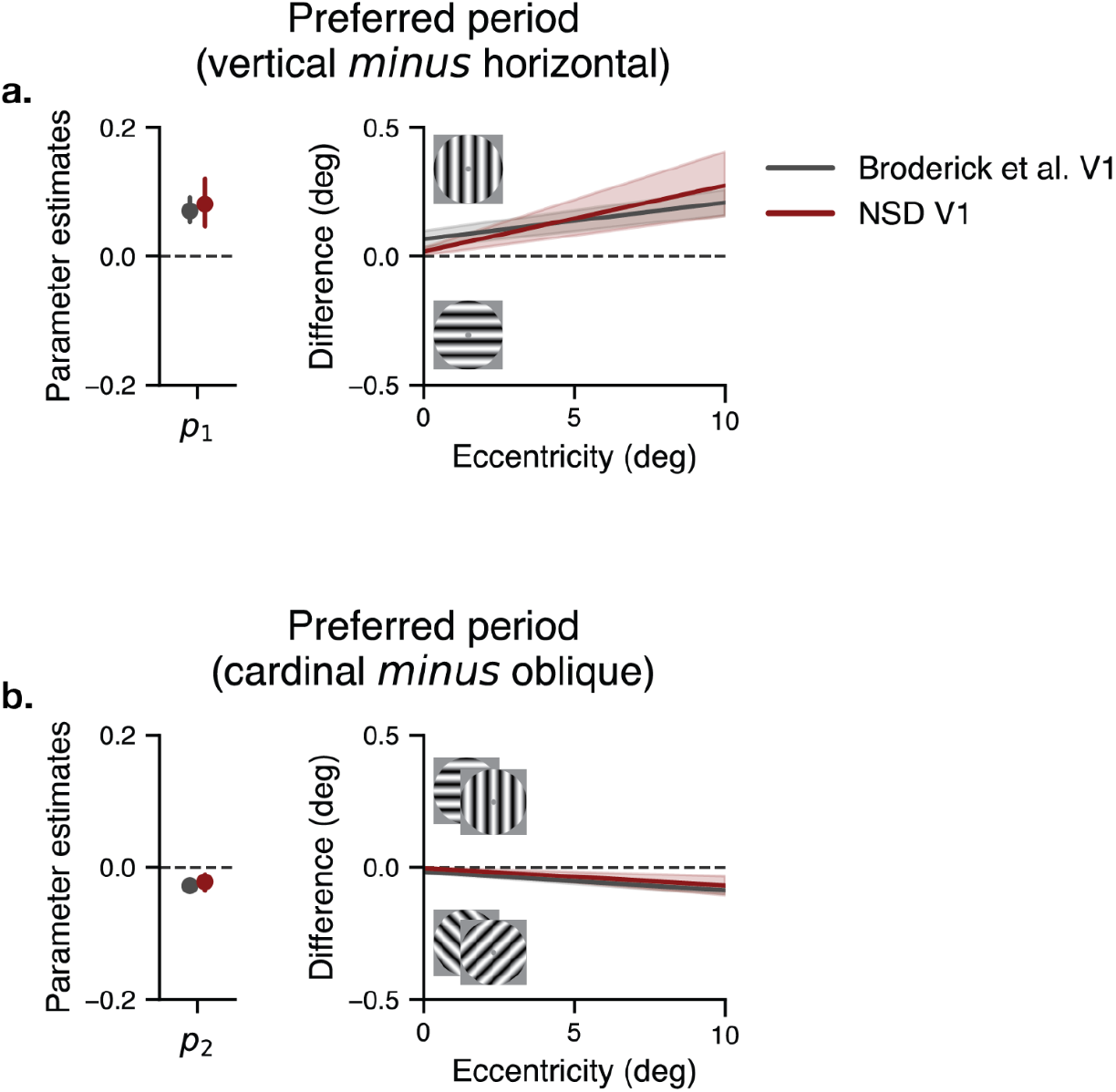
Absolute orientation effect on preferred period of V1 in Broderick et al. and NSD (“Replication”). For all the panels a and b, the left column shows the estimated parameter values. The circles and the error bars represent precision-weighted averages and 68% bootstrapped confidence intervals across subjects. The right column shows the difference in preferred period at local orientation 90° and retinotopic angle 180°. The difference was calculated between the two orientations on which each *p* parameter depends. (a) shows *p*_1_ value in the left panel, and the difference in preferred period between vertical and horizontal orientations in the right panel. (b) shows *p*_2_ in the left panel, and the difference in preferred period between cardinal and oblique orientations in the right panel.

#### 3.2.4. Both datasets show higher BOLD amplitude for vertical than horizontal in V1

The parameters discussed so far pertained to spatial frequency tuning in V1, independent of amplitude. Here we consider the effect of stimulus orientation on amplitude, independent of period. A small but consistent orientation effect on the BOLD responses is observed in both of the datasets. Specifically, in both Broderick et al. and NSD, the *A*_1_ parameter is moderately positive, indicating higher BOLD amplitude for vertical than horizontal orientations (**Figure 9 left**). The two parameter values are about the same, Broderick at 0.04 (0.036, 0.053 68% CI) and NSD at 0.06 (0.046, 0.066 68% CI), meaning that the stimulus orientation modulates the preferred period up or down by 5 or 6%.

**Figure 9.**
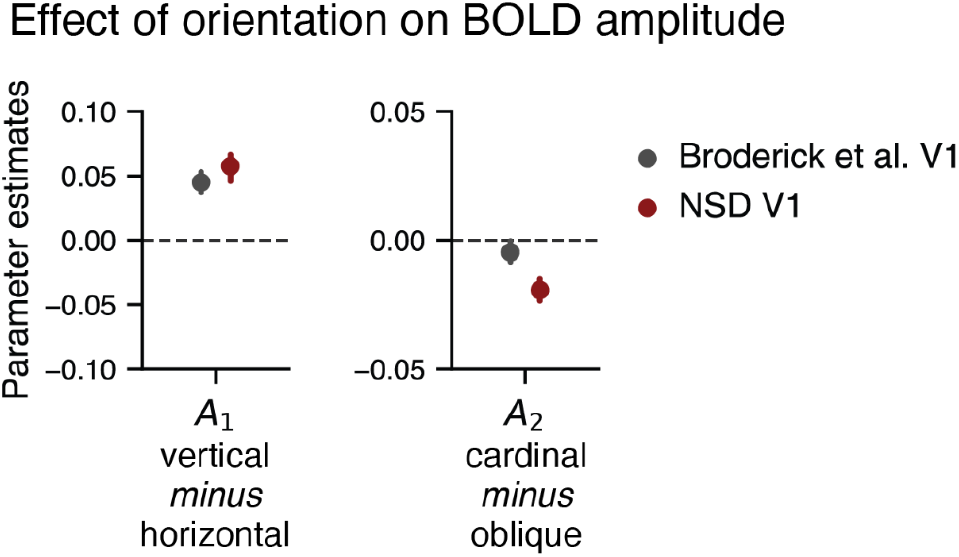
Absolute orientation effect of V1 on BOLD amplitude in Broderick et al. and NSD (“Replication”). The panel shows two estimated parameters (*A*_1,_ *A*_2_). The circles and the error bars represent precision-weighted averages and 68% bootstrapped confidence intervals across subjects.

The two datasets diverge in their estimates for the parameter *A*_2_ (**Figure 9 right**). NSD has a negative *A*_2_, meaning higher amplitude for oblique than cardinal orientations whereas the estimate from Broderick et al. is slightly negative but close to 0. The NSD result aligns with two previous studies reporting stronger responses to obliques (Mannion et al., 2010; Swisher et al., 2010). In contrast, Furmanski and Engel (2000) reported stronger responses to cardinal than oblique orientations, opposite to what we find. However, they only measured on the horizontal meridian, so their finding of greater BOLD amplitude for cardinal than oblique might reflect a radial bias.

#### 3.2.5. Inconsistent effects of relative orientation on preferred period

The last two model parameters are *p*_3_ and *p*_4_, corresponding to orientation relative to retinotopic angle. There were no consistent effects for these parameters. Broderick et al. estimated a positive *p*_3_ value, meaning preferred period was higher for annuli than pinwheels, but NSD did not (**Figure 10a**). The difference might be explained by an interaction with eccentricity. The NSD coverage was more foveal (0.5 to 4.2 deg) than Broderick et al. (1 to 12 deg) and in another study (Himmelberg et al., 2025), the difference in spatial frequency tuning for pinwheels vs annuli reversed with eccentricity: Near the fovea, the peak period was higher for radial patterns, but at 3 deg and beyond, the peak period was higher for annuli.

**Figure 10.**
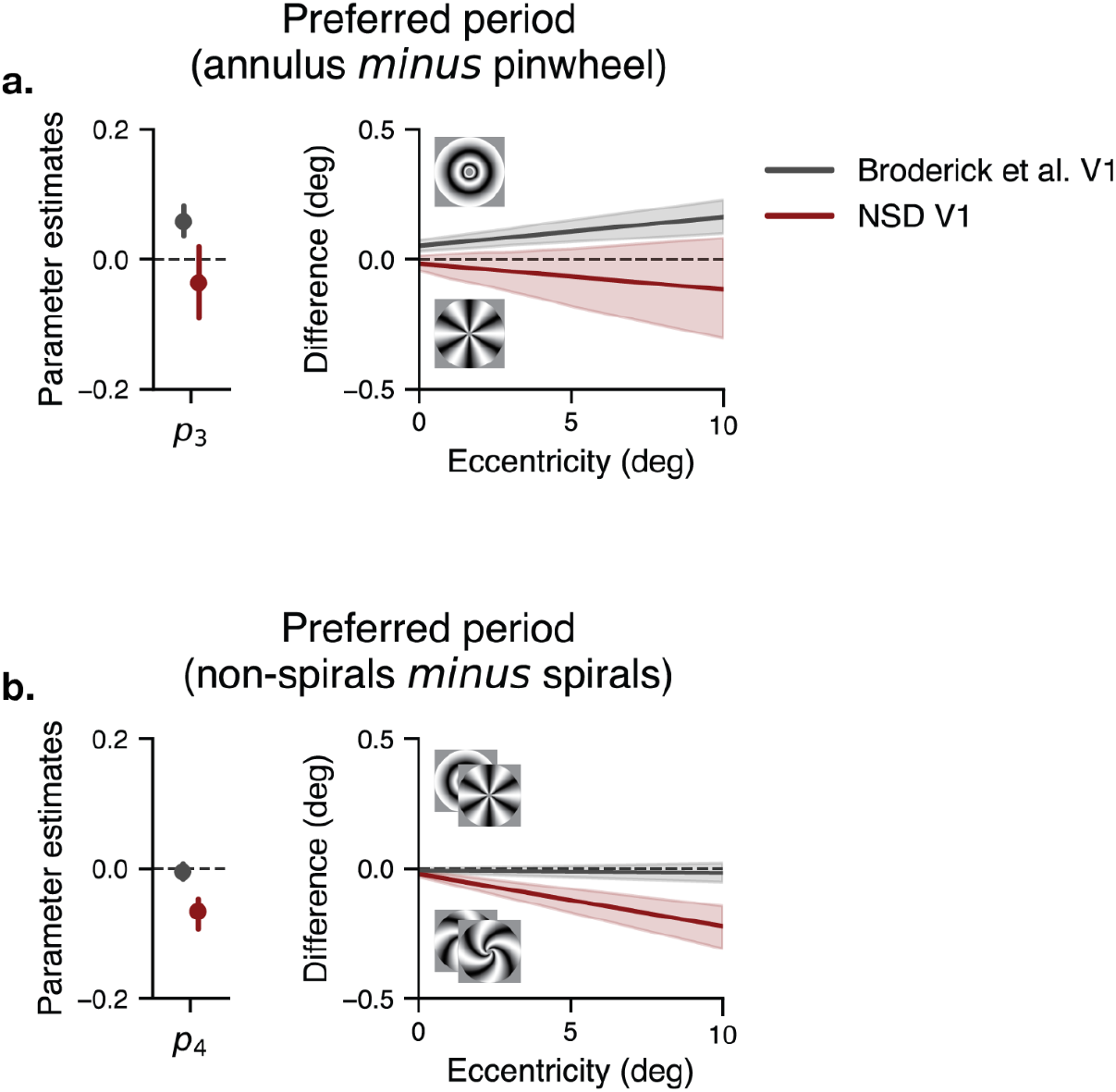
Relative orientation effect on preferred period of V1 in Broderick et al. and NSD (“Replication”). For all the panels a and b, the left column shows the estimated parameter values, whereas the right column shows the difference in preferred period at local orientation at 90° and retinotopic angle at 180°. The difference was calculated between the two orientations on which each *p* parameter depends. In the left column panels, the parameters are labeled after the orientation with a greater preferred period when the values are negative. (a) The results of *p*_3_ which depends on circular (“annulus”) versus radial (“pinwheel”) orientation. (b) The results of *p*_4_ that depends on non-spiral (“annulus” and “pinwheel”) versus spiral (“forward spiral” and “reverse spiral”) orientation. The dots and the error bars represent precision-weighted means and 68% bootstrapped confidence intervals across subjects.

The parameter *p*_4_ was negative in NSD but 0 in Broderick et al. (**Figure 10b**). A negative *p*_4_ indicates a higher preferred period for spirals than non-spirals (annuli or pinwheels).

#### 3.2.6. Replication summary

In sum, the 2D model of V1 spatial frequency tuning in the NSD synthetic dataset reproduced several, though not all, of the effects reported previously by Broderick et al. A summary of all 9 parameters from the two datasets is plotted in **Figure 11**. The bandwidth and effects of eccentricity (especially the combined effect of slope and intercept) were similar in the two datasets. The orientation effects were more complex, but can be summarized in this way: the three orientation parameters with the least cross-participant variability in Broderick et al. were similar between the two studies, whereas the three parameters with the most cross-participant variability in Broderick et al. tended to differ between studies.

**Figure 11.**
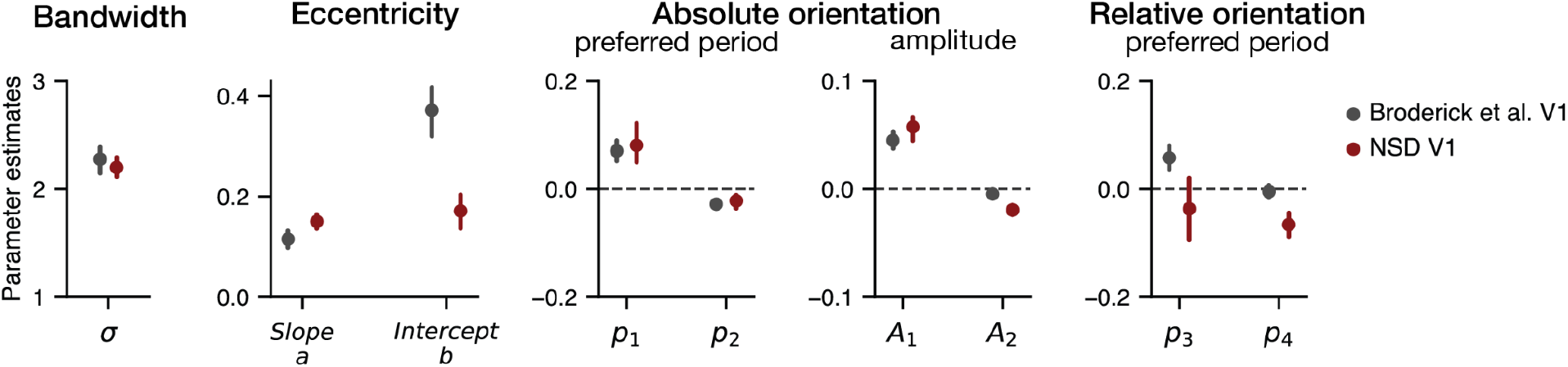
2D parameter estimates for NSD and Broderick et al. for V1. See Supplementary Table 4 for means and confidence intervals of all parameters in both datasets.

Specifically, both studies reliably found higher preferred period for vertical and oblique compared to horizontal and cardinal (*p*_1_ and *p*_2_), and higher amplitude for vertical (*A*_1_), with the mean parameter estimates similar between the two studies. These three parameters had high reliability in Broderick et al, indicated by an effect size (absolute value of mean divided by standard deviation, or Cohen’s *d*) greater than 1: *d*_p1_=1.1, *d*_p2_=1.3, *d*_A1_=1.6.

Among the other three parameters, two differed in that there were reliable effects in NSD (*p*_4_ and *A*_2_) but not Broderick et al. The other parameter, *p*_3_ differed in that it was positive in Broderick (greater preferred period for tangential than radial) and close to 0 in NSD. These three parameters had lower reliability in Broderick et al, with effect sizes lower than 1: *d*_p3_=0.77, *d*_p4_=0.13, *d*_A2_=0.32.

To assess the similarity between the two sets of parameters as a whole, we asked how closely the profile of parameters matched. Since the model parameters vary in units, we first normalized the estimates by their reliability, converting raw values into effect sizes (mean divided by pooled standard deviation). We found that the profile of effect sizes was highly similar between NSD and Broderick et al. (mean squared error=0.69), indicating that the pattern of parameter influence was preserved (**Supplementary Figure 6**). This metric, mean squared error of effect sizes, parallels the ‘replicability in effect size’ used in meta-scientific studies (Open Science Collaboration, 2015). When the analysis was repeated with a null distribution of NSD parameters, the MSE was much larger, from 2-fold to 10-fold across the 1,000 parameter sets from the null distribution (**Supplementary Figures 6, 7)**.

#### 3.2.7. Simulations and parameter recovery

The comparison between the results from the two datasets show that with similar stimuli, one obtains generally similar results. But such agreement does not assess whether the results are correct. It is possible, for example, that the use of stimuli in which the local spatial frequency varied systematically with eccentricity biased the parameter estimates in both studies. To test this possibility, we conducted parameter recovery analyses from two types of simulations.

In the first simulation, we assumed no variation in preferred period as a function of eccentricity, stimulus orientation, or polar angle. This was implemented by setting the slope *m* and all the *p*_n_ terms (effects of orientation on preferred period) to zero. We then generated noisy model responses to scaled gratings and re-estimated the model parameters from the simulated data. If the choice of stimuli introduced bias, the model would erroneously estimate increasing preferred period with eccentricity. Instead, we found that even with a high noise level (3x the measured noise), the model accurately recovered uniform tuning. This result indicates that the relationship between eccentricity and spatial frequency in the stimuli does not, on its own, bias the model toward finding an eccentricity-dependent change in preferred spatial frequency (**Supplementary Figure 2a**).

In a second simulation, we assumed that the parameter values matched the average parameters estimated here (**Figure 11**). The procedure was identical to the first simulation, except that we generated noisy model responses for both scaled gratings and uniform gratings. The resulting parameter estimates were highly similar between the two stimulus sets, indicating that the use of scaled gratings does not impose a large bias in parameter estimates.

### 3.3. The extension of 2D model analysis to V2 and V3

We now extend our analysis by applying the two-dimensional model to extrastriate areas V2 and V3 from NSD. By comparing the 9 parameter estimates across the visual areas, our goal is to identify systematic changes in spatial frequency preferences along the visual hierarchy. To preview the results, we find that the bandwidth of spatial frequency tuning increases from V1 to extrastriate areas, whereas the preferred period does not differ much. Moreover, the oblique effects observed in V1 persist in V2 and V3. Specifically, both the preferred period and the amplitude are higher for oblique orientations than for cardinal orientations in V2 and V3.

#### 3.3.1. Increasing bandwidth from V1 to extrastriate areas

There was a large increase in bandwidth from V1 to V2 and an additional but smaller increase from V2 to V3. The bandwidth parameter *σ* was 2.2±0.1 octaves for V1, 3.8±0.4 octaves for V2, and 4.5±0.3 octaves for V3 (**Figure 12 left**). This corresponds to a full-width-at-half-max of 5.2, 9.0, and 10.6 (V1, V2, and V3). This indicates a 70% increase in bandwidth from V1 to V2, and 100% increase from V1 to V3. To show that the increases in bandwidth from V1 to V2 and from V2 to V3 are reliable within subjects, we plot the parameter differences in the middle panel of **Figure 12**. These differences are large relative to the confidence interval of the differences. Model tuning curves are visualized in the right panel of **Figure 12** across the three areas.

**Figure 12.**
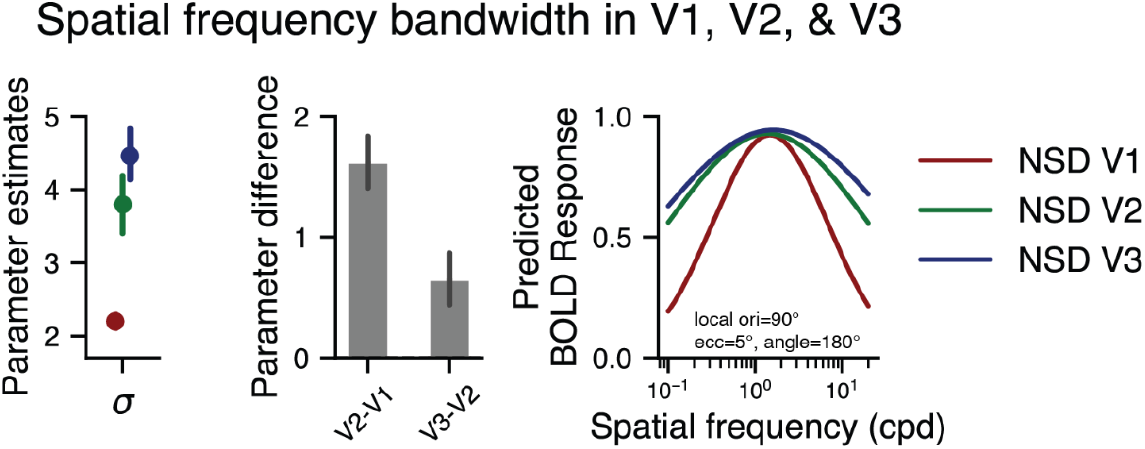
Estimated bandwidth values (left) and spatial frequency tuning curves (right) of V1, V2 and V3 in NSD (“Extension”). NSD V1 data from figure 6 is replotted as a reference. The circles represent precision-weighted averages of bandwidth, and the error bars indicate 68% bootstrapped confidence intervals across subjects. The middle panel shows within-subject differences of the parameter along visual hierarchy. The right panel shows the example model responses, averaged across subjects, assuming local orientation at 90° and a voxel’s pRF eccentricity and angle at 5° and 180°, respectively.

#### 3.3.2. Preferred period does not vary systematically across the V1 to V3 maps

In contrast to the effect of bandwidth, spatial frequency tuning as a function of eccentricity is similar across the V1 to V3 maps. Within each map, the line fit to preferred period vs eccentricity has a positive y-intercept (0.17 to 0.25 deg per cycle at the fovea), and a positive slope (0.15 to 0.17) (**Figure 13 left**). The lines do not show a systematic increase in period with visual areas, with V3 numerically between V1 and V2, though the differences between areas are not large or robust.

**Figure 13.**
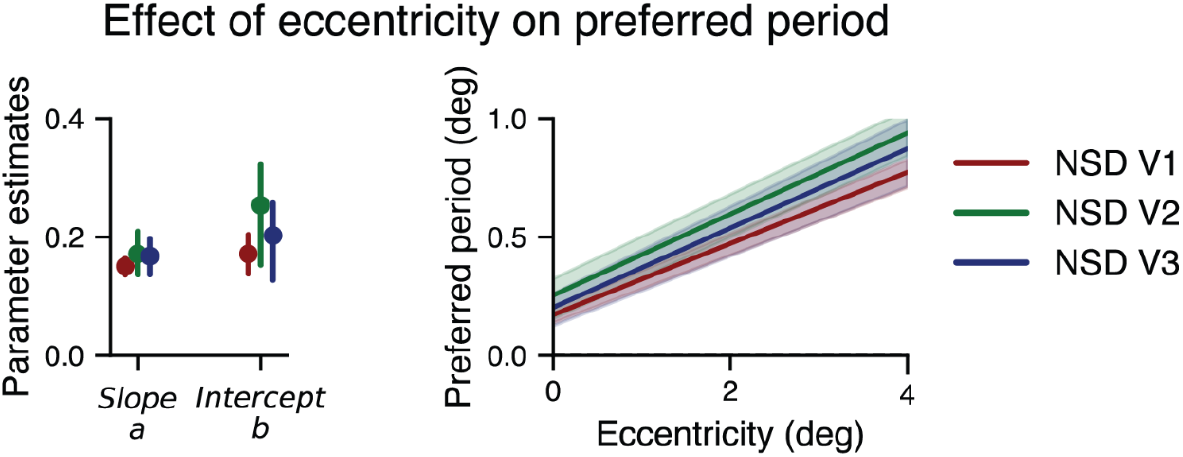
Eccentricity effect on preferred period of V1, V2, and V3 in NSD gratings dataset, with V1 data from Figure 7 replotted as a reference. The left panel shows the estimated slope and intercept values from the two-dimensional model. The circles represent precision-weighted averages. The error bars in both left and right panels indicate 68% bootstrapped confidence intervals across subjects. The right panel shows the predicted preferred period as a function of eccentricity based on the values shown left, with local orientation at 90° and a voxel’s retinotopic angle at 180°.

#### 3.3.3. All three maps have higher preferred period for vertical than horizontal and for oblique than cardinal orientations

In V1, preferred period was slightly but reliably higher for vertical than horizontal orientations. The same pattern was observed in V2 and V3, although the effect is weak in V3. These patterns are summarized by the *p*_1_ parameter estimates, which was 0.08±0.03 for V1, 0.16±0.04 for V2, and 0.05±0.04 for V3 (**Figure 14a**). Additionally, all the early visual areas showed negative values for parameter *p*_2_, indicating that preferred period is higher for oblique than cardinal orientations (**Figure 14b**). Both of these two orientation effects are largest in V2.

**Figure 14.**
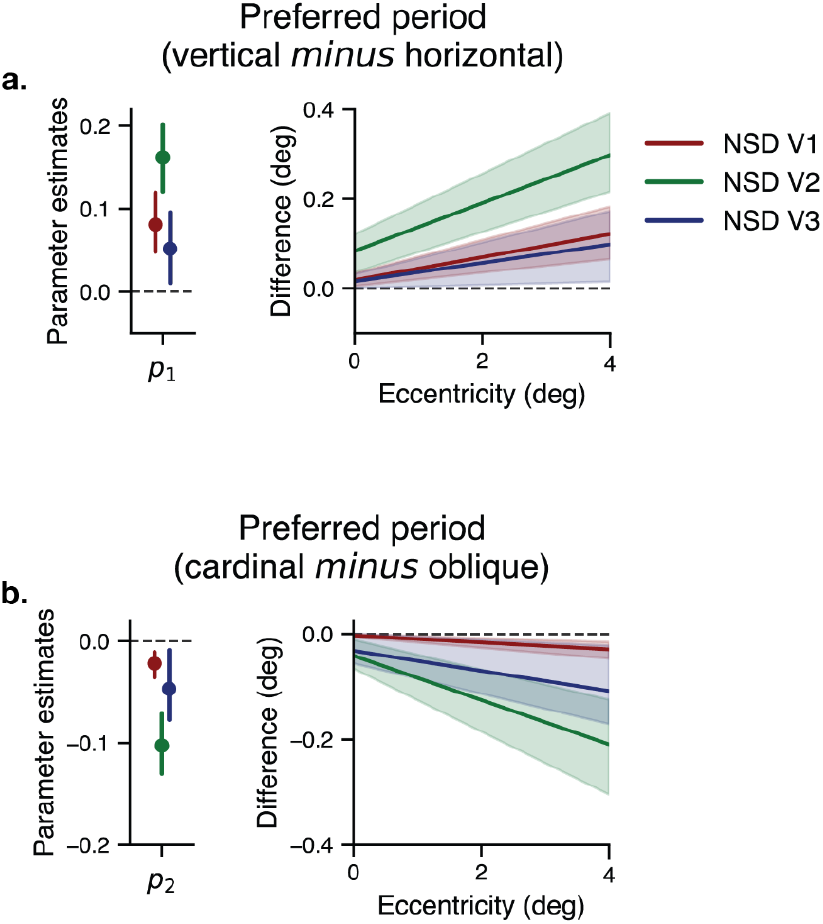
Absolute orientation effect on preferred period of V1, V2 and V3 in NSD (“Extension”), with NSD V1 data from Figure 8 replotted. For all the panels a and b, the left column shows the estimated parameter values. The circles and the error bars represent precision-weighted averages and 68% bootstrapped confidence intervals across subjects. The right column shows the difference in preferred period at local orientation 90° and retinotopic angle 180°. The difference was calculated between the two orientations on which each *p* parameter depends. (a) shows the difference in preferred period between vertical and horizontal orientations. (b) shows the difference in preferred period between cardinal and oblique orientations.

#### 3.3.4. Effects of absolute orientation on amplitude are similar across V1, V2, and V3

Next, we examined the effect of stimulus orientation of extrastriate areas on BOLD amplitude. The parameter *A*_1_, which contrasts amplitude for vertical vs horizontal, was positive for V1, V2, and V3 (**Figure 15**). The parameter *A*_2_, which measures amplitude for oblique compared to cardinal, was negative for all three maps, indicating higher amplitude for oblique than cardinal orientations. Thus, in all three maps in NSD, for oblique stimuli, the preferred period is higher (*p*_2_) and the amplitude is higher (*A*_2_).

**Figure 15.**
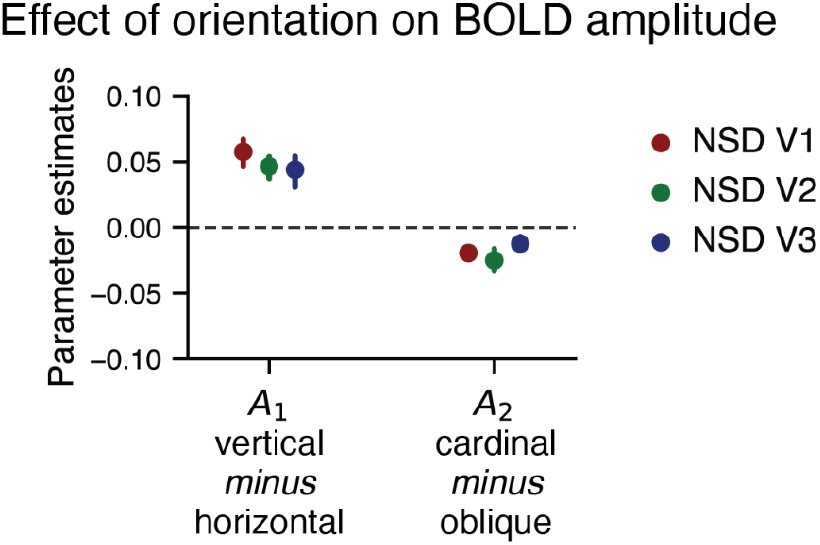
Absolute orientation effect on BOLD amplitude in NSD V1, V2, and V3 (“extension”). The panel shows two estimated parameters (*A*_1,_ *A*_2_) for each visual area. The circles and the error bars represent precision-weighted averages and 68% bootstrapped confidence intervals across subjects.

Note that all of the amplitude effects, even when statistically reliable, are small: for example, the *A*_2_ parameter for V2 is about -0.02±0.01, meaning that the predicted BOLD responses to oblique gratings and cardinal gratings differ by about 4%. More concretely, if the mean BOLD response in a voxel were, say, 1% signal change, then the model would predict 1.02% for oblique and 0.98% for cardinal.

#### 3.3.5. No systematic effects of relative orientation on preferred period across V1, V2 and V3

As described in section 3.2.5, the two parameters for relative orientation, *p*_3_ and *p*_4_ were not consistent between Broderick et al. and NSD for V1. Here we see that the parameters also show no systematic effects across V1, V2, and V3 in NSD. In NSD V1, *p*_3_ is about 0 (no difference in preferred period for radial vs circular patterns) and *p*_4_ is negative (higher preferred period for spirals than non-spirals). In contrast, the V2 and V3 estimates are negative for *p*_3_ and about 0 for *p*_4_, but there is a lot of variability across subjects in the V2/V3 estimates of *p*_3_ and *p*_4,_ making it difficult to determine whether the differences from V1 are meaningful (**Figure 16**).

**Figure 16.**
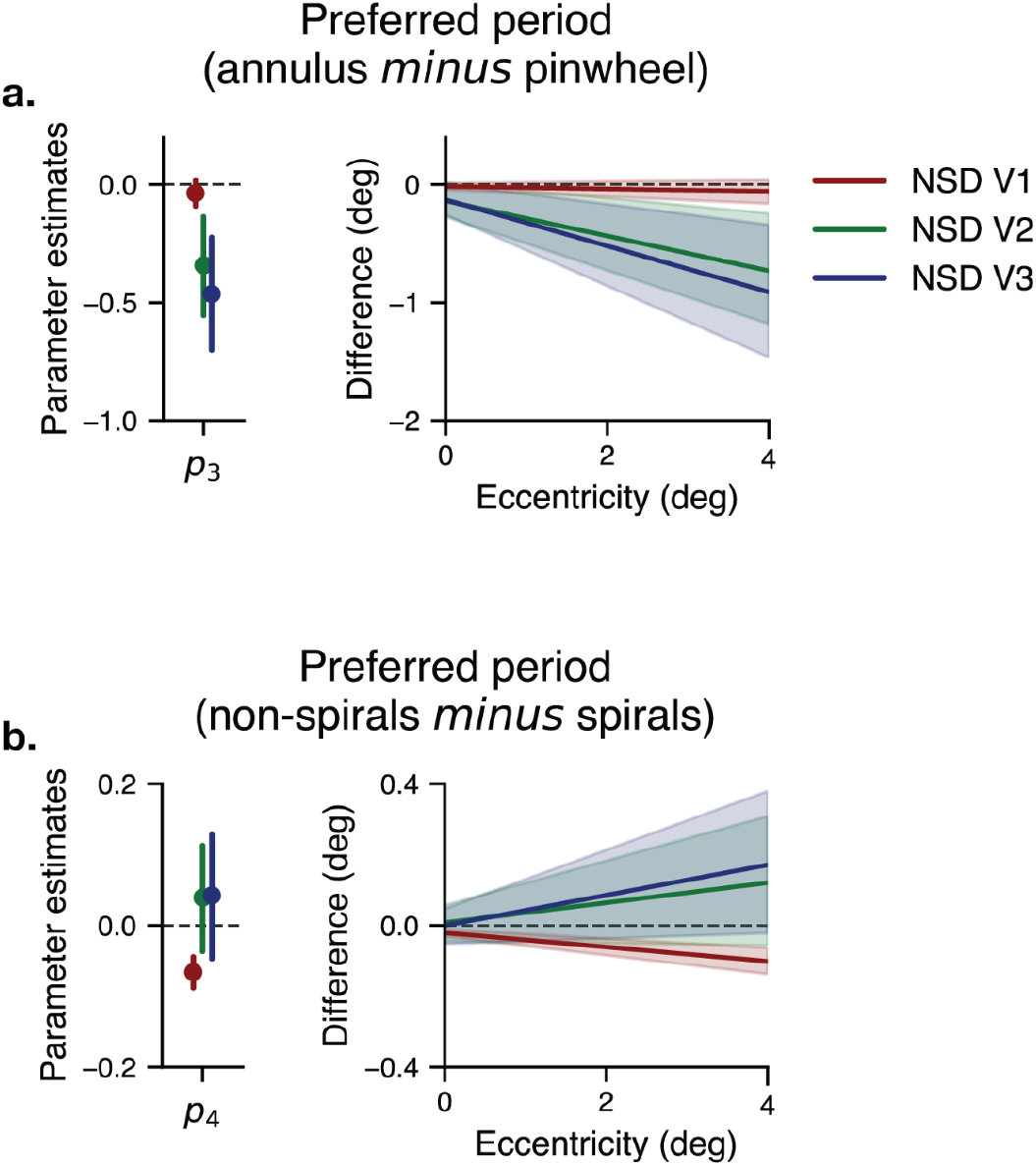
Relative orientation effect on preferred period of V2 and V3 in NSD (“Extension”), with NSD V1 from Figure 10 replotted. For all the panels a and b, the left column shows the estimated parameter values, with the dots and the error bars representing precision-weighted averages and 68% bootstrapped confidence intervals across subjects. The right column shows the difference in preferred period when local orientation is at 90° and retinotopic angle is at 180°. The difference was calculated between the two orientations on which each *p* parameter depends. (a) The difference in preferred period between circular (“annulus”) and radial (“pinwheel”) orientations. (b) The difference in preferred period between non-spiral (“annulus” and “pinwheel”) and spiral (“forward spiral” and “reverse spiral”) orientations.

#### 3.3.6. Extension summary

First, we observed that in NSD, there were reliable responses to our stimuli in V2 and V3, with clear spatial frequency tuning. This enabled us to quantify the effect of eccentricity on peak spatial frequency tuning, which was similar to the effects in V1. The most salient difference between the extrastriate areas and V1 was the increase in bandwidth. Moreover, the model fits show similar effects of absolute orientation in the extrastriate maps in V1– higher preferred period and higher amplitude for vertical than horizontal stimuli (*p*_1_, *A*_1_) and for oblique than cardinal stimuli (*p*_2_, *A*_2_). The effects of relative orientation were less systematic (*p*_3_, *p*_4_). A summary of the 9 parameters for all three visual areas in NSD is plotted in **Figure 17**.

**Figure 17.**
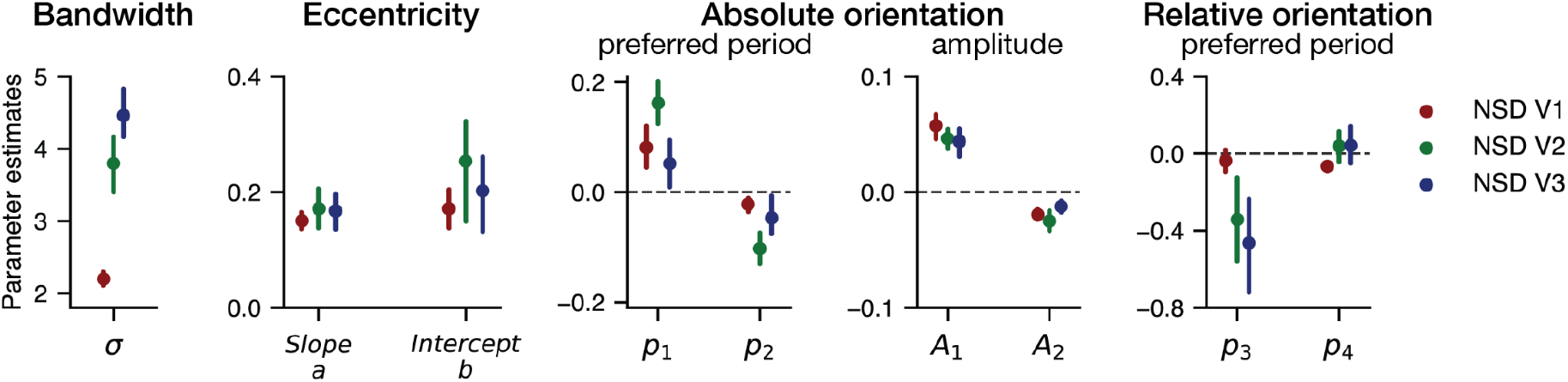
2D model fits for all the nine parameters of NSD V1, V2, V3. For means and confidence intervals of all parameters in each area, see Supplementary Table 4.

## 4. Discussion

We measured and modeled spatial frequency tuning in human visual cortex using data from the NSD. When implementing the same 9-parameter model of V1 spatial frequency tuning as described in Broderick et al. (2022), the profile of parameters was similar between the two studies. In particular, the bandwidth was nearly identical, and the preferred frequency as a function of eccentricity was also closely matched, albeit with a lower intercept and slightly steeper slope in the NSD. Three of the six orientation parameters were also highly similar, whereas three were not. The orientation parameters that were similar were those that showed more consistent effects across subjects in the original study (Cohen’s *d* > 1). The three that did replicate had lower consistency in the original study (Cohen’s *d* < 1).

We then applied the model to V2 and V3 data from NSD, and compared the model parameters to those for V1. The most salient difference between the extrastriate maps and V1 was an increase in bandwidth, doubling from V1 to V3.

### 4.1. Computational reproducibility

In order to assess whether the results from Broderick et al replicate, it is essential to show *computational reproducibility*, that the same results are obtained from a re-analysis of the original data. Otherwise, any differences in parameters from the previously reported parameters would be ambiguous, as differences could arise from either differences in the datasets or the algorithm. The data and code from Broderick et al. were both publicly available (http://hdl.handle.net/2451/63344), as were the data from NSD synthetic, facilitating a test of computational reproducibility, and indeed the re-analysis showed computational reproducibility.

In addition to *whether* results are computationally reproducible, an important issue is *the ease* of reproducibility. It has been argued that “frictionless reproducibility” is the critical factor needed for rapid scientific progress (Donoho, 2024; Recht, 2024). In this case, the process was not frictionless: an error was discovered in the original report in the stimulus description (Broderick et al., 2024) and refitting the models required optimizing hyperparameters and other model-fitting procedures, consistent with the challenges intrinsic to reproducing neuroscience results (Wandell, 2024) and to the sensitivity of neural analyses on processing pipelines (Botvinik-Nezer et al., 2020; Li et al., 2024).

### 4.2. Empirical reproducibility

*Empirical reproducibility* refers to whether one obtains the same results from a new study. The 2D model used to analyze the data was identical to that used by Broderick et al., and the scaled grating stimuli were highly similar, although there were also several differences, including magnetic field strength, fMRI acquisition and processing, experimental design, and stimulus size (Table 1). Many of the results from Broderick et al. were similar in the NSD dataset, despite the differences between the experiments. In particular, the V1 bandwidth and the effects of eccentricity were quite similar between the studies. Three recent papers (including this one) have converged to close agreement on peak spatial frequency tuning as a function of eccentricity in V1 (Aghajari et al., 2020; Broderick et al., 2022), greatly narrowing the uncertainty from prior studies (D’Souza et al. 2016; Farivar et al. 2017; Henriksson et al. 2008; Sasaki et al. 2001; Hess et al. 2009; Kay 2011) (**Supplemental Figure 4**).

The effects of orientation on spatial frequency are much smaller than those of eccentricity. For example, the preferred period more than doubles from 1 deg to 4 deg, but varies by only 10% as a function of orientation. It is therefore perhaps not surprising that these effects do not replicate as well as the eccentricity effects. Nonetheless, the three orientation parameters with relatively high consistency across subjects in Broderick et al. (Cohen’s *d* > 1) led to very similar results in NSD, increasing confidence that these effects, although small, are robust and generalizable.

The similarity of our V1 model parameters to those of Broderick et al. provides some assurance that our V1 model is reasonable, and justifies our comparison of the V1 results to those of extrastriate cortex in NSD.

### 4.3. Similar preferred period along visual hierarchy

The most salient finding from the extrastriate data is that bandwidth increases, doubling from V1 to V3, whereas preferred period remains consistent. The relatively consistent preferred period across areas might seem at odds with the notion that V1 neurons, with small receptive fields, code fine details, whereas higher visual areas, with larger receptive fields, are sensitive to structure at a larger spatial scale. But in fact, larger receptive fields need not imply a preference for lower spatial frequencies. If, for instance, a V2 neuron pools from many V1 neurons with different spatial frequency tuning and different receptive field locations, the V2 neuron’s preferred frequency will be similar to the average preferred frequency of its V1 afferent, but its receptive field size will be larger, matching the combined receptive fields of its afferent. Hence the V2 neuron’s size will be larger than those in V1 but its preferred frequency will be similar. Consistent with this idea, behavioral judgments that are presumed to rely on mid-level or high-level visual representations can be driven by high spatial-frequency content (Lieber et al., 2023). Such results suggest that the increasingly large receptive fields in higher visual areas are not necessarily associated with a shift in preference to lower spatial frequencies, consistent with our findings in V1 to V3.

### 4.4. Increasing bandwidth along the visual hierarchy

The increasing bandwidth from V1 to V3 parallels patterns observed with spatial receptive fields: in extrastriate visual maps, receptive fields are larger at corresponding map locations. Might the patterns with spatial frequency tuning and spatial receptive fields arise from the same cause? Consider a simple pooling model in which the relationship between V1 and V2 receptive field properties arise from convergent feedforward connectivity. Assuming V2 neurons pool over local populations of V1 neurons, then V2 receptive field centers will match the receptive field center averaged across the V1 population providing inputs (i.e., similar centers), whereas the V2 receptive field size will match the aggregate size of that population (i.e. larger receptive fields). Harvey and Dumoulin (2011) applied this logic to infer the spatial extent of V1 that each V2 location pools over: they found that with a small Gaussian pooling function (3-mm radius) applied to the V1 surface, they could predict V2 pRF size from V1 properties (pRF size and the cortical magnification function).

To assess pooling effects, we simulated the effects of applying a 3-mm Gaussian pooling to V1 outputs (**Supplementary Analysis 1**). This results in a 65% increase in receptive field size, comparable to the increase in size from V1 to V2. Surprisingly, the same pooling function for spatial frequency tuning predicted less than 1% increase in spatial frequency bandwidth, inconsistent with the 70% increase in bandwidth we observed in the data **(Supplementary Figure 5)**. Hence pooling over just a few mm of cortex substantially increases spatial receptive field size but has little effect on spatial frequency bandwidth. The reason is that position tuning changes rapidly along the V1 map, whereas the preferred period changes slowly. An additional explanation is needed.

One likely explanation is that the greater bandwidth in extrastriate areas arises from nonlinearities in addition to pooling. V2 shows increasing non-linearities in spatial tuning (Kay et al. 2013), temporal tuning (Zhou et al., 2018), and contrast sensitivity (Kay et al. 2013) compared to V1. The increasingly sharp nonlinearities along the visual hierarchy can arise from a canonical nonlinearity computed at each processing stage (Kay et al. 2013). A similar architecture is employed in hierarchical neural network models of the visual system (Riesenhuber *&* Poggio, 1999). Such nonlinearities can broaden tuning or increase invariance to stimulus features. Consider a high spatial frequency pattern that only weakly drives V1 neurons in a patch of cortex. V2 neurons that pool the responses of V1 neurons nonlinearly might amplify the weak response, effectively increasing the V2 bandwidth. In our simulation, pooling followed by a compressive power-law nonlinearity (square root) results in 42% broader tuning. Smaller exponents would result in even larger increases.

A possible alternative explanation is that there are a relatively small number of V1 neurons tuned to very low frequencies or very high frequencies, with minimal effect on the V1 BOLD signal, and that V2 oversamples these outputs. We cannot distinguish the two possibilities with the current data and analyses. However, successive stages of pooling followed by compressive nonlinearities is a more parsimonious explanation, as it can account for the increasing non-linearities observed across multiple stimulus domains, e.g., both space and time (Kim et al., 2024).

### 4.5. Relation to prior studies

Our finding of increasing bandwidth but constant preferred period from V1 to V2 to V3 differs from some prior fMRI studies. Aghajari et al. (2020) found the opposite pattern: similar bandwidths and decreasing preferred frequency from V1 to V3. Henriksson et al. (2008) also found decreasing preferred spatial frequency from V1 to V2, though like this study, they did not find an orderly progression from V1 to V2 to V3: they found that preferred spatial frequency was similar for V1 and V3 at low eccentricities, and for V2 and V3 at higher eccentricities. The differences between our results and those of Aghajari et al. ‘s arise from the extrastriate data, not V1. For example, at 5 deg eccentricity, our bandwidth estimate for V1 is similar to theirs (5.2 vs 4.5 octaves), whereas ours increase in V2 and V3 (9.0 and 10.6 octaves) and theirs do not. The differences could arise from differences in experimental design or analysis. We suspect that experimental design is most important.

The experiments differed in both the stimulus sequence and the stimulus content. We used a randomized, event-related design and Aghajari et al used a periodic sweep of spatial frequencies. The spatial frequency sweep, like the traveling wave in retinotopy (Engel et al., 1994), affords a higher signal-to-noise ratio. But it also can cause a large bias in bandwidth estimate due to incorrect assumptions about the hemodynamic response function, similar to the error in size estimation that can occur with pRF fitting (Lerma-Usabiaga et al., 2020). The event-related design, though lower in SNR, is not susceptible to this bias. If the assumed hRF differs from the true hRF more in extrastriate areas than V1, this could explain the observed discrepancies between studies. In support of this interpretation, a recent study measuring spatial frequency tuning with an event-related design also showed increasing bandwidth from V1 to V2 to V3 (Subramanian et al., 2026), though a smaller increase than what we find in NSD (50% increase from V1 to V3 compared to doubling). Subramanian et al. also observed only a slight decrease in preferred spatial frequency from V1 to V3, comparable to our observations from NSD.

The stimulus content also differed between studies, with Aghajari et al. using band-pass noise patterns whose spatial frequency does not vary with eccentricity, whereas Broderick et al. and NSD using scaled gratings. These differences might matter most for extrastriate areas. Response selectivity in V1 is reasonably well explained by models assuming an underlying linear filter (e.g., Heeger, 1992). In contrast, phase and higher order properties of stimuli can modulate spatial frequency tuning in high-level visual cortex (Berman et al., 2017). More generally, neural encoding in extrastriate visual areas is sensitive to higher order image statistics, including second order contrast (Freeman et al., 2013; Movshon *&* Simoncelli, 2014; Okazawa et al., 2017), and as a result, measurements of spatial frequency tuning in these areas are more likely to depend on image properties beyond the local spatial frequency. Data from Henriksson et al. (2008) is consistent with this interpretation. They found that the preferred spatial frequency in V2 increased as angular sinusoidal patterns grew in length (from 90º to 180º to 360º), whereas the preferred frequency in V1 changed little.

Our study and Aghajari et al.’s also differed in analysis method, with Aghajari et al. fitting voxel-wise tuning curves allowing for different bandwidths for each voxel, whereas we fit a single model to all the data within a map, assuming a single bandwidth. The much larger number of parameters in the voxel-wise approach makes the estimation of each parameter more susceptible to noise. In contrast, imposing a low-dimensional model on an entire visual map reduces the susceptibility to noise, but at the cost of possible increase in bias (i.e., bias-variance tradeoff, Hastie, 2009). There are several advantages to the area-based approach, as discussed by Broderick et al. But the difference in analysis method alone is unlikely to explain the discrepancy in results, since our results are in good agreement for V1, differing only in V2 and V3.

One factor not included in the 9-parameter model used in this paper and by Broderick et al. is the effect of polar angle. In fact, preferred spatial frequency in V1 does appear to vary systematically with polar angle (Aghajari et al., 2020; Himmelberg et al., 2025). Preferred spatial frequency is about 30% higher on the horizontal the vertical meridian, and about 17% higher on the lower than upper vertical meridian. The effect of polar angle is much smaller than eccentricity, but it nonetheless indicates that spatial frequency tuning varies along multiple retinotopic dimensions.

### 4.6. Opposing effects on amplitude and preferred spatial frequency

A curious finding is that stimulus orientations that have higher preferred spatial frequency elicit lower amplitude responses. This was evident in the *p*_1_ and *A*_1_ parameters (vertical > horizontal) and the *p*_2_ and *A*_2_ parameters (oblique > cardinal). In other words, for horizontal orientations compared to vertical, the preferred spatial frequency is higher and the amplitude is lower. And similarly, for cardinal orientations compared to oblique, the preferred spatial frequency is higher and the amplitude is lower. These patterns were generally consistent, in that they were found in both Broderick et al. and NSD V1, and in NSD V2 and V3. One might expect that tuning to higher frequencies would support better visual acuity. Indeed, both grating acuity (Berkley et al., 1975) and vernier acuity (Corwin et al., 1977; Leibowitz, 1955) are higher for cardinal than oblique orientations, paralleling our finding that spatial frequency tuning is higher for cardinal stimuli. There is not, however, clear evidence that acuity is higher for horizontal than vertical orientations (Berkley et al., 1975; Campbell et al., 1966; Long *&* Tuck, 1991). Thus the link between the cortical tuning measured by fMRI and behavior is not straightforward. Moreover, higher BOLD amplitude for oblique than cardinal orientation is not straightforward to link to poorer performance on most tasks with oblique orientations than cardinal orientations, as stimuli that elicit higher amplitude neural responses are often associated with better behavioral performance, at least on detection tasks (Ress et al., 2000).

More generally, there are many challenges in linking behavioral performance to neural data. One is that the neural measures, as in this paper, are made with high contrast, large stimuli to elicit large responses, whereas behavioral measures are often made with low contrast, small stimuli to measure local thresholds. Second, without designing the behavioral and neural studies together, there are likely to be differences in stimuli (duration, eccentricity, spatial extent, and so on), subject populations, and other methods. And third, perhaps most importantly, a neural circuit model of the task is needed in order to know how the neural measure should limit performance (Kay et al., 2023). Such models are difficult to implement for most kinds of experiments (Morgan et al., 2013). Although we do not offer linking models in this paper, we hope that the data we report, including the measures of variability between subjects and replicability across subjects, can serve as useful constraints in the development of such models.

## 5. Conclusions

Three reports, including this one, now show quantitative agreement on the spatial frequency tuning in human V1 as a function of eccentricity (Aghajari et al., 2020; Broderick et al., 2022), despite differences in stimuli, analysis methods, and scanners. This is an advance over where the field stood just a few years earlier, where the range of reported values for peak spatial frequency tuning in V1 varied several-fold across studies. Much of the advance is likely due to a combination of stimulus selection, spanning a wide range of spatial frequencies, and improvements in analysis methods. The increased precision in the characterization of spatial frequency tuning has further enabled replicable measures of more subtle effects. In particular, effects of absolute stimulus orientation on spatial frequency tuning also replicated across studies. Finally, the most novel observation reported here is that the bandwidth of spatial frequency tuning increased substantially from V1 to V2 and V3, suggesting a parallel between position tuning and spatial frequency tuning: both receptive field size and spatial frequency bandwidth increase along the cortical visual hierarchy.

## 7. Data and Code Availability

The data and script necessary to replicate them are available via the Open Science Framework (https://osf.io/umqkw) and Github (https://github.com/JiyeongHa/Spatial-Frequency-Preferences_NSDsyn).

## 8. Author Contributions

Ha conducted all the analyses with input from Broderick and Winawer. Ha drafted the manuscript with input from Winawer. All authors edited the manuscript. Kay was responsible for NSD data collection and provided guidance on accessing and analyzing the NSD data. Winawer oversaw the project.

