## Supplementary information for "Spatial Frequency Maps in Human Visual Cortex: A Replication and Extension"

#### Supplementary Table 1. Frequency information of experimental stimuli.

**Spatial frequency properties of the scaled grating stimuli.** For each stimulus class, the table shows the radial frequency ($\omega_r$), angular frequency ($\omega_a$), and the resulting base frequency ($\omega$) along with an example local spatial frequency ($\omega_l$) at 2° eccentricity.

| Stimulus Class | $\omega_r$ | $\omega_a$ | $\omega$ | $\omega_{l=2º}$ |
|---|---|---|---|---|
| Pinwheels | 0 | 6 | 6 | 3 |
| | 0 | 11 | 11 | 5.5 |
| | 0 | 20 | 20 | 10 |
| | 0 | 37 | 37 | 18.5 |
| | 0 | 69 | 69 | 34.5 |
| | 0 | 128 | 128 | 64 |
| Annuli | 6 | 0 | 6 | 3 |
| | 11 | 0 | 11 | 5.5 |
| | 20 | 0 | 20 | 10 |
| | 37 | 0 | 37 | 18.5 |
| | 69 | 0 | 69 | 34.5 |
| | 128 | 0 | 128 | 64 |
| Forward spirals | 4 | 4 | 5.66 | 2.83 |
| | 7 | 7 | 9.90 | 4.95 |
| | 14 | 14 | 19.80 | 9.90 |
| | 26 | 26 | 36.76 | 18.38 |
| | 49 | 49 | 69.29 | 34.64 |
| | 91 | 91 | 128.67 | 64.34 |
| Reverse spirals | 4 | -4 | 5.66 | 2.83 |
| | 7 | -7 | 9.90 | 4.95 |
| | 14 | -14 | 19.80 | 9.90 |
| | 26 | -26 | 36.76 | 18.38 |
| | 49 | -49 | 69.29 | 34.64 |
| | 91 | -91 | 128.67 | 64.34 |
| Intermediate spirals | 14 | 34 | 36.78 | 18.39 |
| | 34 | 14 | 36.78 | 18.39 |
| | 34 | -14 | 36.78 | 18.39 |
| | 14 | -34 | 36.78 | 18.39 |

#### Supplementary Table 2. Difference in MRI acquisition parameters between the Broderick et al. (2022) and NSD gratings dataset.

| | Broderick et al. (2022) dataset | NSD gratings dataset |
|---|---|---|
| MR field strength | 3T | 7T |
| Voxel resolution | 2 mm iso | 1.8 mm iso |
| Coverage | 208 mm x 208 mm inplane, 66 slices | 216 mm x 216 mm inplane, 84 slices |
| Pulse sequence | Grad echo EPI (CMRR Multi-Band EPI sequences) | Grad echo EPI (CMRR Multi-Band EPI sequences) |
| TR, TE, flip angle | 1000 ms; 37 ms; 68° | 1,600 ms; 22 ms; 62° |
| Sample rate | 1 s | 1.6 s |
| Phase encoding direction | Poster-to-Anterior | Anterior-to-Posterior |
| Acceleration | multi-band slice acceleration factor 6 | Partial Fourier 7/8, iPAT 2, multi-band slice acceleration factor 3 |
| Coil | Siemens 64 channel head and neck RF coil | Siemens 32 channel RF head coil |
| Distortion correction | Dual echo field maps | Dual echo field maps |
| Task | fixation-related | fixation-related or image-related |

#### Supplementary Table 3. NSD synthetic vs Broderick et al. dataset functional MR image preprocessing pipeline

| Preprocessing steps | NSD synthetic (Gifford et al., 2025) | Broderick et al. (2022) |
|---|---|---|
| Motion correction | SPM5 *spm_realign* | FSL *mcflirt* |
| Slice timing correction | Temporal resampling with cubic interpolation; upsampled to 1.0s (1mm) TR | None |
| Distortion correction | Time-varying fieldmaps (phase–unwrapped using FSL utility *prelude*, regularization using 5mm Epanechnikov kernel, linearly interpolated) | Reversed phase-encoding scans (FSL TOPUP / ApplyTOPUP) |
| Registration with anatomical space | Mean fMRI volume (1 mm, averaged across 5 NSD sessions) co-registered to T2 (ANTs *BSplineSyN*), allowed a small amount of nonlinearity | Co-registered to T1w using boundary-based registration with 9 degrees of freedom (Freesurfer *bbregister*) |
| Spatial smoothing | None | None |
| Temporal filtering | None | None |
| Temporal resampling | Cubic interpolation to upsample the data (1.0s) | None |
| GLM estimation | 1 β per trial (*GLMsingle*) | 1 β per condition (*GLMdenoise*) |
| Surface reconstruction | Freesurfer *recon-all* | Freesurfer *recon-all* |
| Step order | Temporal resampling → Distortion correction → Motion correction → Registration → Surface mapping | Motion correction → Distortion correction → Registration → Surface mapping |

#### Supplementary Table 4. Precision-weighted averages and bootstrapped confidence intervals for the 2D model across datasets and ROIs.

**Parameter estimates and with bootstrapped confidential intervals.** For each parameter (rows) and dataset-ROI group (columns), values show the precision-weighted mean followed by 68% confidence interval [lower, upper] and 95% confidence interval [lower, upper] in brackets. Confidence intervals were estimated using 1,000 bootstrap resamples with percentile method.

| parameter | Broderick et al. V1 | NSD V1 | NSD V2 | NSD V3 |
|---|---|---|---|---|
| sigma | 2.28<br>[2.16, 2.38]<br>[2.07, 2.54] | 2.2<br>[2.11, 2.29]<br>[2.03, 2.38] | 3.8<br>[3.41, 4.19]<br>[3.23, 4.6] | 4.46<br>[4.17, 4.83]<br>[3.89, 5.21] |
| slope | 0.12<br>[0.1, 0.13]<br>[0.08, 0.15] | 0.15<br>[0.14, 0.16]<br>[0.12, 0.18] | 0.17<br>[0.14, 0.21]<br>[0.12, 0.25] | 0.17<br>[0.14, 0.19]<br>[0.1, 0.22] |
| intercept | 0.37<br>[0.32, 0.42]<br>[0.29, 0.47] | 0.17<br>[0.14, 0.2]<br>[0.1, 0.23] | 0.25<br>[0.15, 0.32]<br>[0.07, 0.39] | 0.2<br>[0.13, 0.26]<br>[0.06, 0.32] |
| $p_1$ | 0.071<br>[0.052, 0.09]<br>[0.036, 0.109] | 0.081<br>[0.048, 0.12]<br>[0.01, 0.134] | 0.162<br>[0.126, 0.2]<br>[0.085, 0.232] | 0.052<br>[0.006, 0.096]<br>[-0.02, 0.147] |
| $p_2$ | -0.028<br>[-0.034, -0.022]<br>[-0.04, -0.017] | -0.022<br>[-0.035, -0.011]<br>[-0.051, -0.003] | -0.103<br>[-0.129, -0.072]<br>[-0.16, -0.049] | -0.047<br>[-0.077, -0.011]<br>[-0.105, 0.012] |
| $p_3$ | 0.058<br>[0.037, 0.08]<br>[0.015, 0.103] | -0.037<br>[-0.094, 0.021]<br>[-0.149, 0.057] | -0.342<br>[-0.567, -0.136]<br>[-0.755, -0.017] | -0.464<br>[-0.717, -0.249]<br>[-0.918, -0.065] |
| $p_4$ | -0.005<br>[-0.016, 0.006]<br>[-0.028, 0.021] | -0.066<br>[-0.09, -0.044]<br>[-0.114, -0.024] | 0.039<br>[-0.043, 0.124]<br>[-0.08, 0.209] | 0.042<br>[-0.05, 0.129]<br>[-0.12, 0.224] |
| $A_1$ | 0.045<br>[0.037, 0.053]<br>[0.03, 0.061] | 0.058<br>[0.046, 0.067]<br>[0.033, 0.074] | 0.046<br>[0.037, 0.054]<br>[0.026, 0.059] | 0.044<br>[0.031, 0.055]<br>[0.021, 0.064] |
| $A_2$ | -0.005<br>[-0.009, -0.0]<br>[-0.013, 0.003] | -0.019<br>[-0.024, -0.015]<br>[-0.028, -0.012] | -0.025<br>[-0.033, -0.017]<br>[-0.041, -0.008] | -0.012<br>[-0.017, -0.007]<br>[-0.021, -0.002] |

#### Supplementary Figure 1. Cross-validation

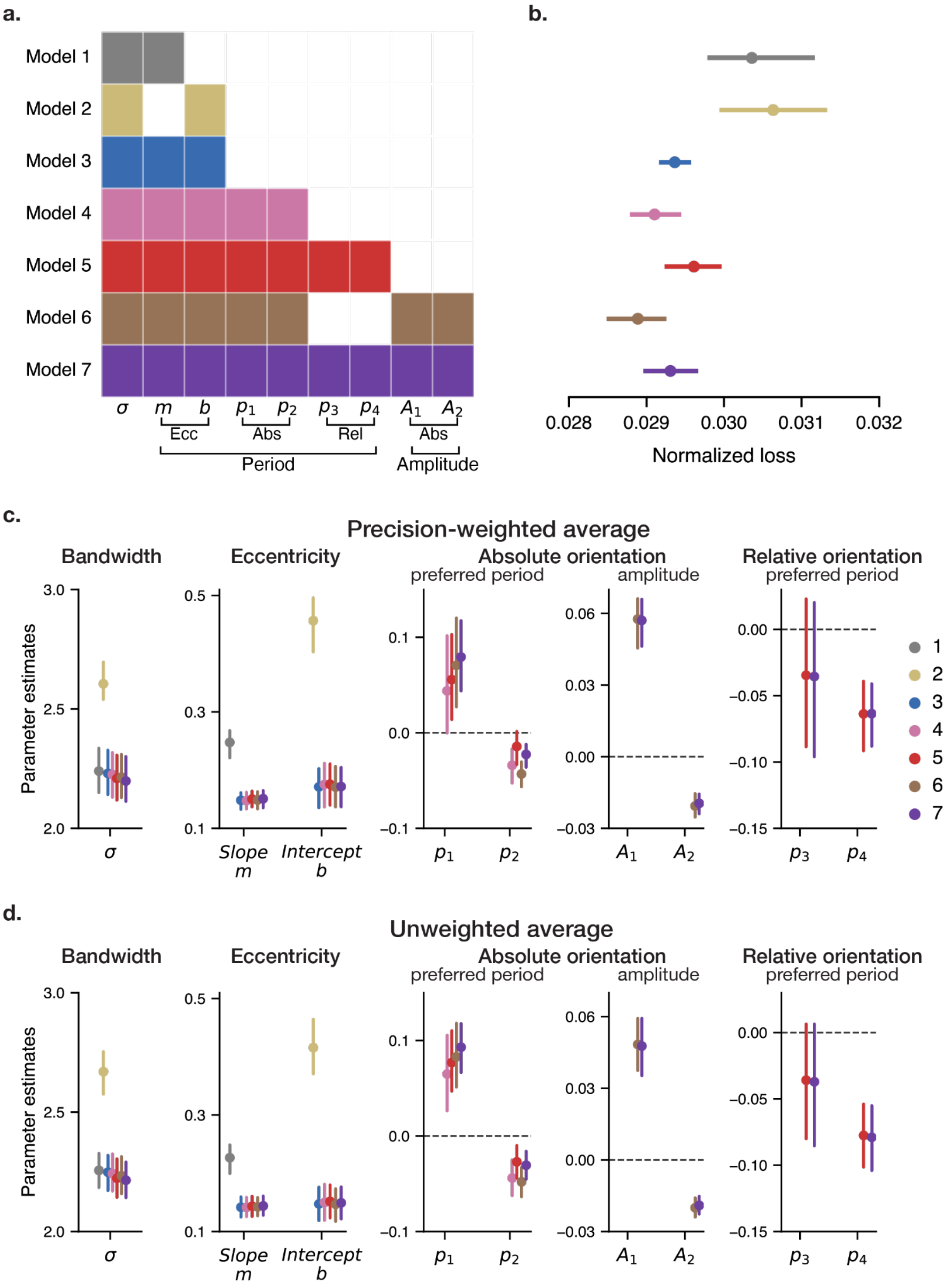


**Model comparison. (a)** Seven models differing in which parameters are included. Model parameters are grouped by whether they affect preferred frequency or amplitude, and whether their effects relate to eccentricity, absolute orientation, or relative orientation. **(b)** Cross-validated loss for each model. Each model is fit to each subject's data using 7-fold cross-validation (train on 24, test on 4). Each subject has one loss per model. The loss is normalized by subtracting the subject-wise mean across models, then re-centered by adding back the grand mean across all subjects and models, as in Broderick et al. Error bars indicate 68% confidence intervals bootstrapped across subjects (1000 bootstraps). **(c)** Parameter estimates for each model fit to the full dataset, weighted by subject precision. Error bars indicate 68% confidence intervals bootstrapped across subjects. **(d)** Same as c, except averaged with equal weights across subjects.

### Supplementary Figure 2. Parameter recovery

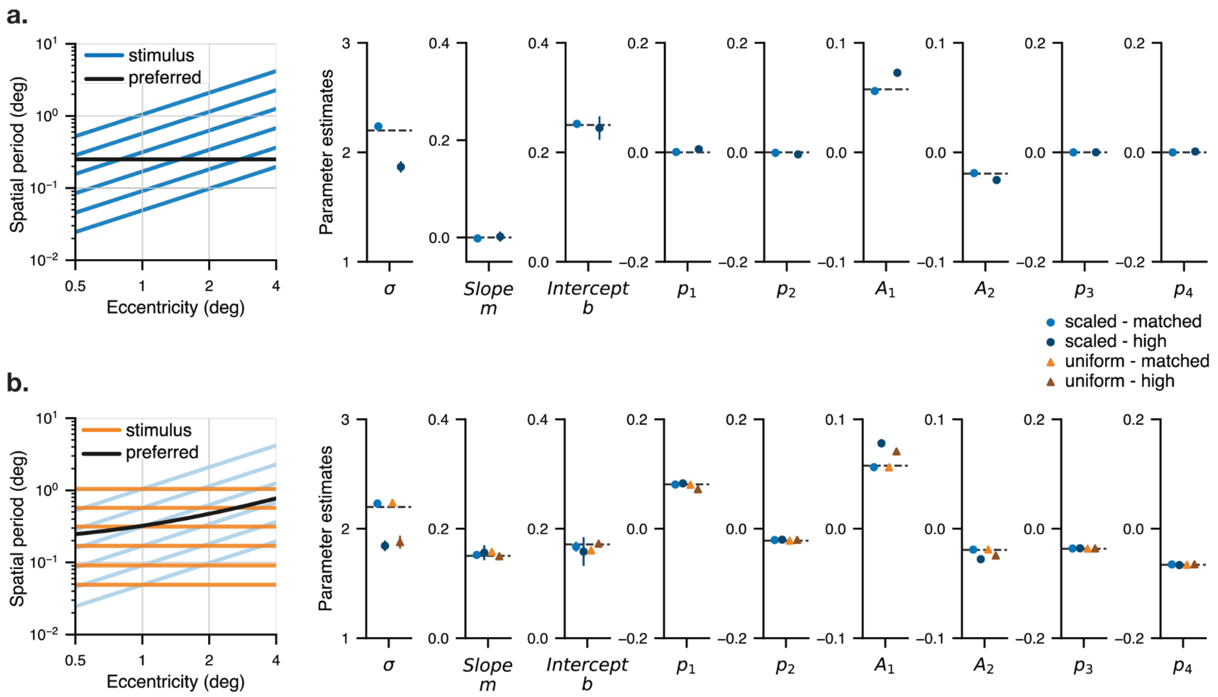


**Simulations for assessing potential stimulus sampling bias. (a)** Simulations in which the ground-truth effects of eccentricity and orientation on spatial period were set to zero. The intercept was set to 0.25, and the other parameters (bandwidth, and the effects of orientation on amplitudes) were set to the precision-weighted average values estimated from this study. The left panel plots the preferred period (black line, flat) and the stimulus period (blue lines) as a function of eccentricity. The blue lines slope upward because the simulations used scaled gratings. Both axes are plotted on a logarithmic scale for better visualization. Two hundred simulations were conducted, 100 with "matched noise" and 100 with "high noise". Error bars (±SD across 100 simulations with different random seeds) are nearly invisible, indicating highly stable parameter estimates. The parameter values for simulations are plotted as dashed black lines (0 for *m* and $p_n$). "Matched" and "high" noise means the data were simulated using a noise covariance matrix that was the same as, or 3x higher than, the noise covariance matrix estimated from subject01's data. The noise covariance in the data was estimated using a toolbox from Generative Modeling of Signal and Noise (GSN; Kay et al. 2025). The results show that with both low noise (light blue dots) and high noise (dark blue dots), the recovered parameters were close to the ground truth, with no evidence that scaled gratings imposed eccentricity or orientation effects. **(b)** Same as panel A, except for two differences. First, the ground truth parameter values were matched to those reported in the present study (averaged across subjects). Second, separate simulations were conducted with uniform gratings (orange lines in left plot) and scaled gratings (light blue lines in left plot). The scaled-grating stimulus periods are identical to those in the left panel (a), but shaded light blue here for clarity. The results show that with both matched and high noise, for both uniform and scaled gratings, parameters were generally recovered accurately. There were some subtle effects of high noise, in that the bandwidth (sigma) was lower than ground truth, but the values did not differ between scaled and uniform gratings.

### Supplementary Figure 3. Model parameter estimates for individual subjects

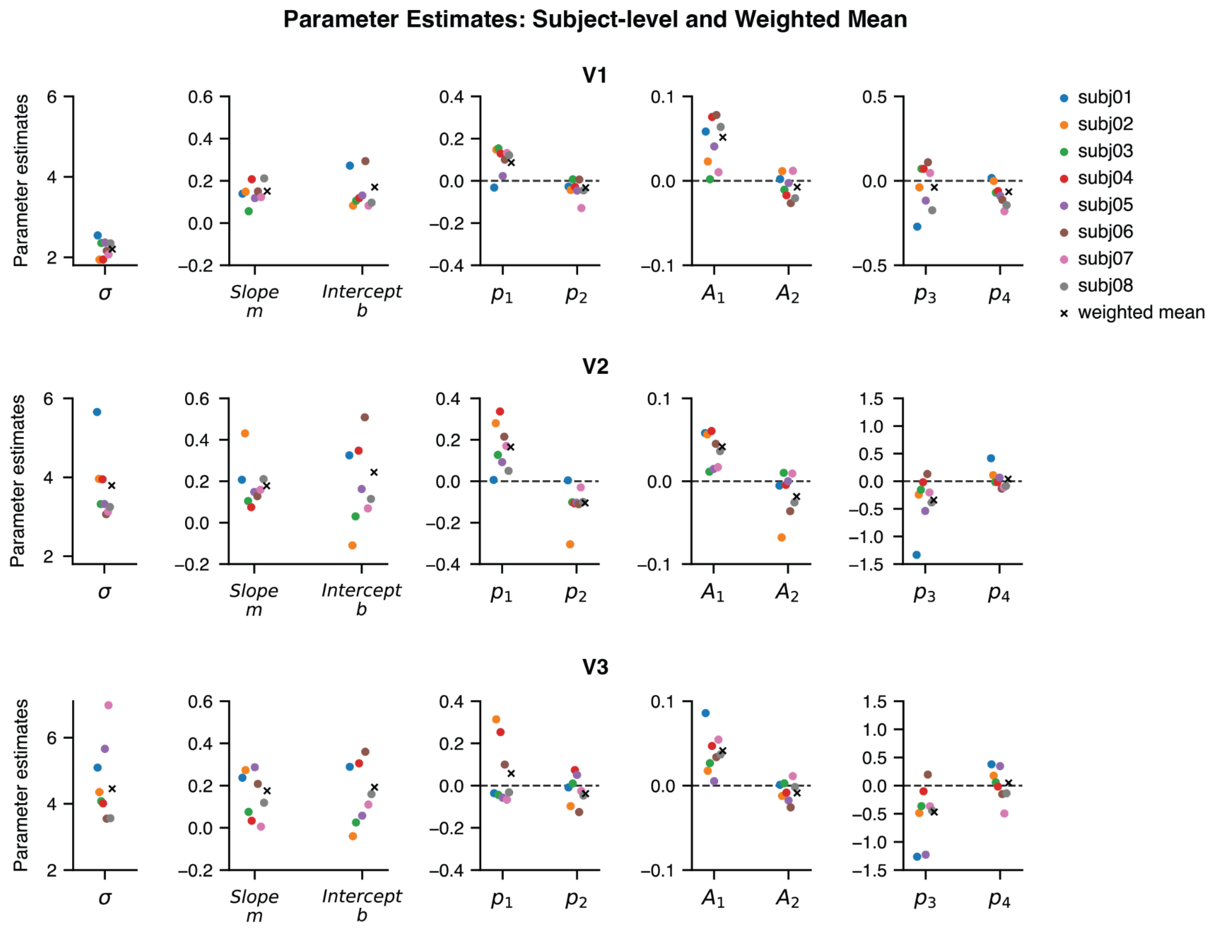


**Subject-level parameter estimates and weighted means.** Parameter estimates in V1, V2, and V3 for individual subjects. Weighted means across subjects reported in this study are replotted for comparison (black crosses). Each point reflects a model fit to data averaged over 8 repeated trials and 4 phases. Further details on the experimental design are provided in Section 2.1.1 (NSD experimental design).

### Supplementary Figure 4. Comparison with previous fMRI studies

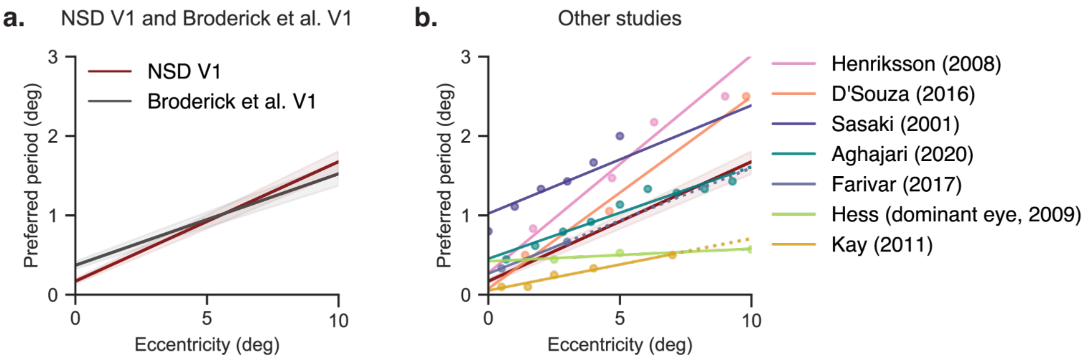


**Spatial period preferences as a function of eccentricity across fMRI studies. (a)** The NSD and Broderick et al. studies (left) have similar preferences within the central 10º (replotted from Figure 7 as precision-weighted averages and 68% bootstrapped confidence intervals across subjects). **(b)** Spatial period preferences from 7 other fMRI studies are plotted for comparison to the NSD, with regression lines (solid) and extrapolation lines (dotted). This panel is adapted from Broderick et al. 2022 (their Figure 11). The preferred period estimates vary 4-fold across studies at the 5º (mid-eccentricity) point.

#### Supplementary Analysis 1. Simulation of population pooling effects

We conducted a simulation analysis to help understand the relationship between spatial frequency tuning in V1 and V2, particularly the large increase in bandwidth we observed in the data. To do so, we made the assumption that V2 neurons pool signals from a local region of V1, and we then compared the properties of the V1 and V2 neural populations. The simulations included effects of preferred position (receptive field center), receptive field size, preferred spatial frequency and spatial frequency bandwidth.

We first simulated a 1D array of V1 population receptive fields (pRFs) along a line from fovea to periphery, spaced every 0.5 mm from 0 mm (fovea) to 40 mm (10 deg eccentricity). Each local population was assigned 4 properties:

1. pRF center (eccentricity, in deg)
2. size (Gaussian SD, in deg)
3. preferred spatial frequency (cycles/deg)
4. bandwidth (octaves).

1. The pRF centers $E$ (in deg), as a function of cortical position $X$ (in mm), were derived from Horton and Hoyt's (1991) cortical magnification function:

$$Magnification = \frac{17.3}{E+0.75},$$

$$E = 0.75 \cdot (e^{\frac{X}{17.3}} - 1)$$

2. The pRF size was determined by the linear function relating size to eccentricity from (Dumoulin and Wandell 2008) (their figure 9):

$$pRF\ size = 0.06E + 0.3$$

3. The preferred spatial period $P$ (in deg/cycle) was defined by the slope $m$ and intercept $b$ parameters in our models (averaged across participants):

$$P = 0.15E + 0.17P$$

4. The bandwidth $\sigma$ (in octaves) was fixed at a single value, as in our 9-parameter model:

$$\sigma = 2.2$$

The 1D profiles of these receptive fields and spatial frequency tuning functions are plotted in **Supplementary Figure 5 (**left panels of 5a and 5b). Each column is one cortical location, with 0.5 mm spacing between them. The colormap within each column shows the tuning curve, either for position (panel a) or spatial period (panel b).

To simulate pooling across a local region of V1, the V1 maps (one for position, one for spatial frequency) were convolved with a Gaussian kernel of 3.2 mm, corresponding to the population point-image size reported by Harvey & Dumoulin (2011). This operation yielded the "pooled" outputs in the middle panels of **Supplementary Figure 5a**. The red and green vertical lines indicate the same arbitrary location in all plots, 25 mm. The tuning curves at this location are plotted in the rightmost column, for spatial location in panel *a* and spatial period in panel *b*. The red curves plot the V1 tuning curves from the left column. The green curves plot the “V2” tuning curves from the middle column, i.e. the V1 tuning curves after spatial pooling. Finally, to approximate post-pooling normalization, the pooled responses were raised to a power-law nonlinearity (exponent $n = 0.5$), generating the normalized pRF profiles, plotted in blue dashed lines.

#### Supplementary Figure 5. Effect of pooling on neural properties

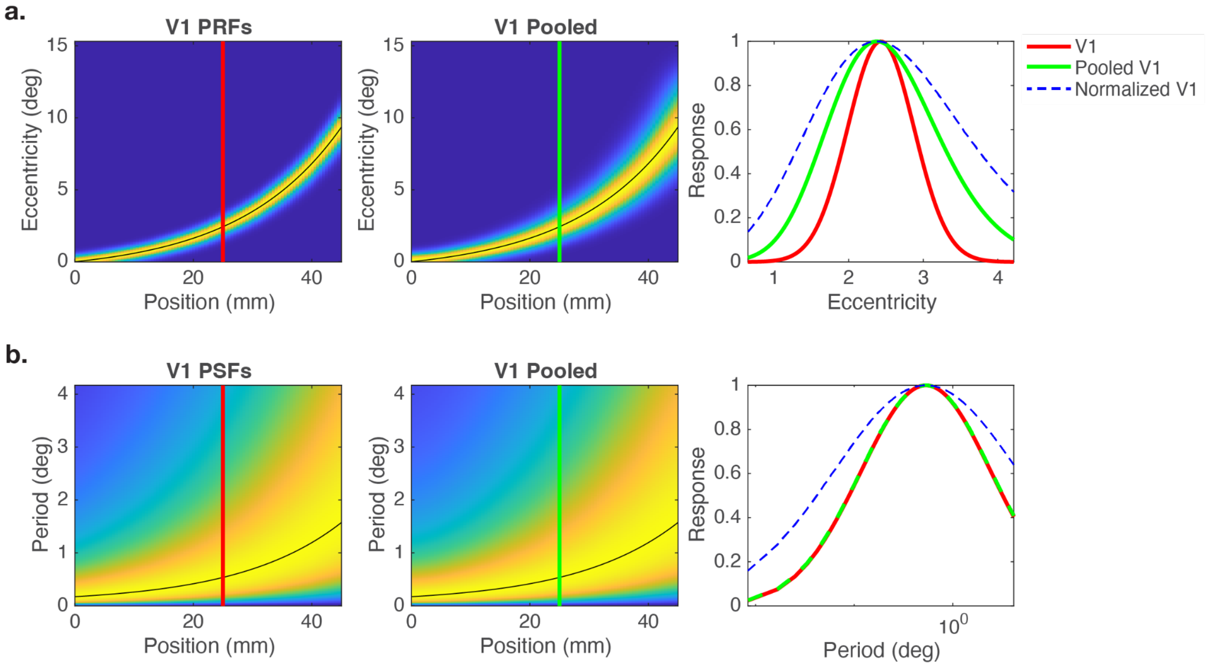


**c.** PRF sizes and PSF sizes from simulation at 25 mm (2.5° eccentricity)

| | V1 | Pooled V1 | Normalized V1 |
|---|---|---|---|
| PRF size (deg) | 0.45 | 0.74 | 1.06 |
| PSF size (octaves) | 2.20 | 2.21 | 3.13 |

**Effect of pooling on neural properties. (a)** A 1D slice of simulated V1 population receptive fields (PRFs), from fovea (0º, 0 mm) to periphery (10º, 45 mm). The columns are spaced every 0.5 mm. The colormap within the columns showed the spatial receptive field of that local population. The black curve traces the preferred location (pRF center) across the map. The middle panel is the same except that the left column has been convolved with a Gaussian kernel (σ = 3.2 mm) to simulate the effects of V2 neural populations pooling over a region of V1. The rightmost column plots example tuning functions from the left two panels corresponding to the red and green vertical lines, that is, the colormaps along a column in the left two panels are converted to a line plot in the right panel. The blue dashed line plots the pooled tuning function after applying a power-law nonlinearity ($n = 0.5$). **(b)** Same as panel a but for spatial period instead of spatial position. **(c)** Widths of the tuning functions plotted in the rightmost panels of A and B. The simulation demonstrates that pooling greatly broadens pRF tuning (65% increase) but has little effect on PSF tuning (<1%). The power law nonlinearity increases bandwidth in both functions.  attributable to the nonlinearity. The figure and table can be reproduced using code located at /figure/suppl_fig5.m at https://github.com/JiyeongHa/Spatial-Frequency-Preferences_NSdsyn/

**Supplementary Figure 6. Similarity of model parameters between NSD dataset and Broderick. et al.**

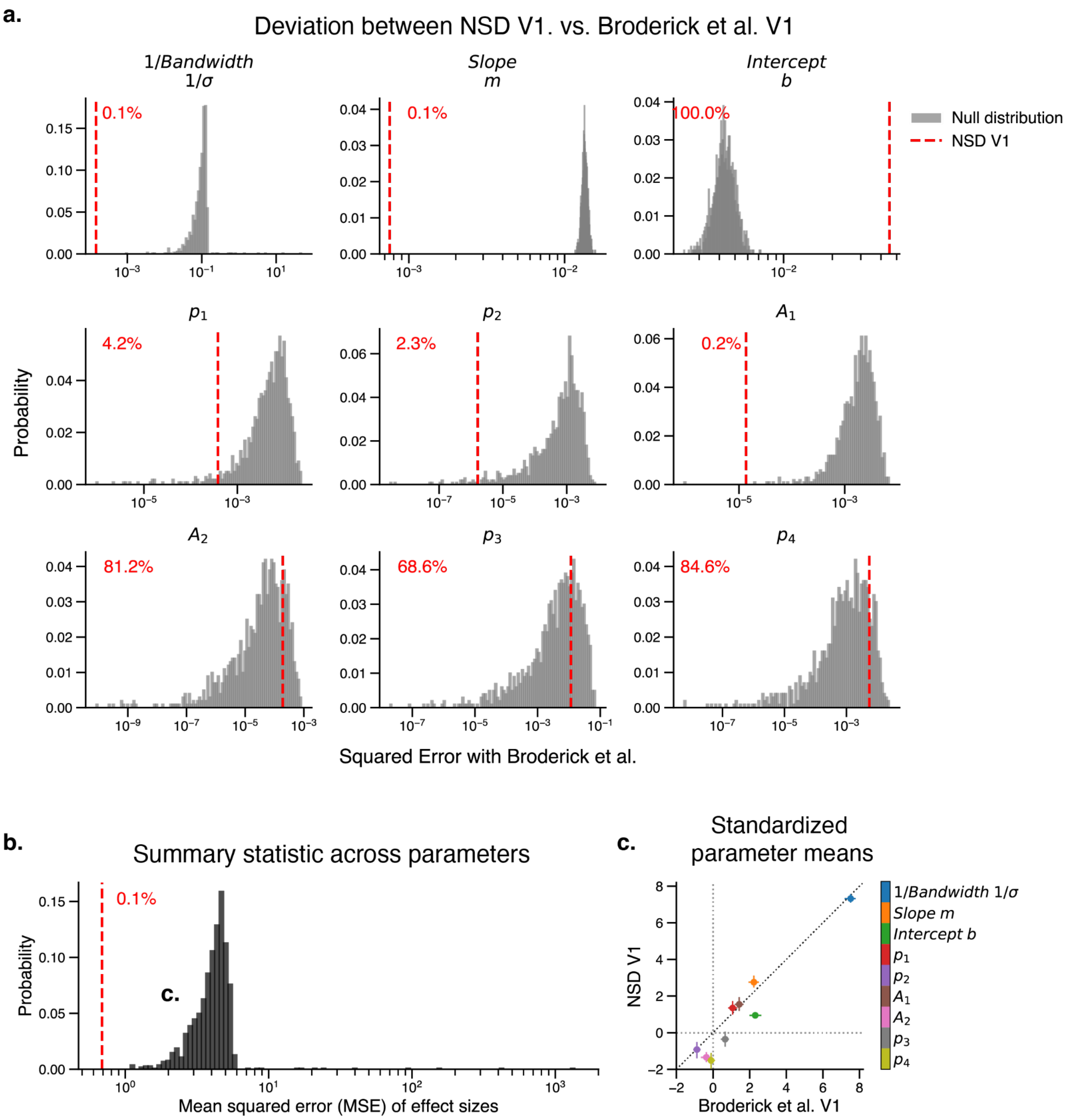


**Parameter similarity between NSD and Broderick et al. V1. (a) Differences between Broderick et al and NSD for each of the 9 parameters.** In each plot, the red dashed line is the squared difference between the means of the two studies, after conversion of each parameter to effect size (mean divided by the pooled standard deviation across subjects). The gray histograms are the squared differences between the mean of Broderick et al, and each of the 1,000 parameters sets from shuffling NSD (see section 2.5 in Methods). The red text is the percentile squared error of the observed data relative to the null distribution. See Supplementary Figure 7 for the corresponding distributions of each parameter in original units (not transformed to effect size and not converted to squared error). **(b) Summary across parameters.** The similarity between the two vectors of 9 parameters was summarized by the mean squared error. As with panel A, the values for the unshuffled NSD are plotted as red dashed lines, and for the null distribution as histograms for the 1,000 bootstrap samples, and the red number indicates the percentile of the unshuffled data relative to the null distribution. The MSE between Broderick et al and NSD without shuffling is lower than the MSE of all 1,000 parameter sets in the shuffled distribution. **(c)** A scatterplot of the 9 pairs of parameter estimates (in units of effect size) for Broderick et al and NSD (unpermuted). The error bars are 68% CIs across participants.

#### Supplementary Figure 7. Null distributions of 2D model parameter estimates in original units

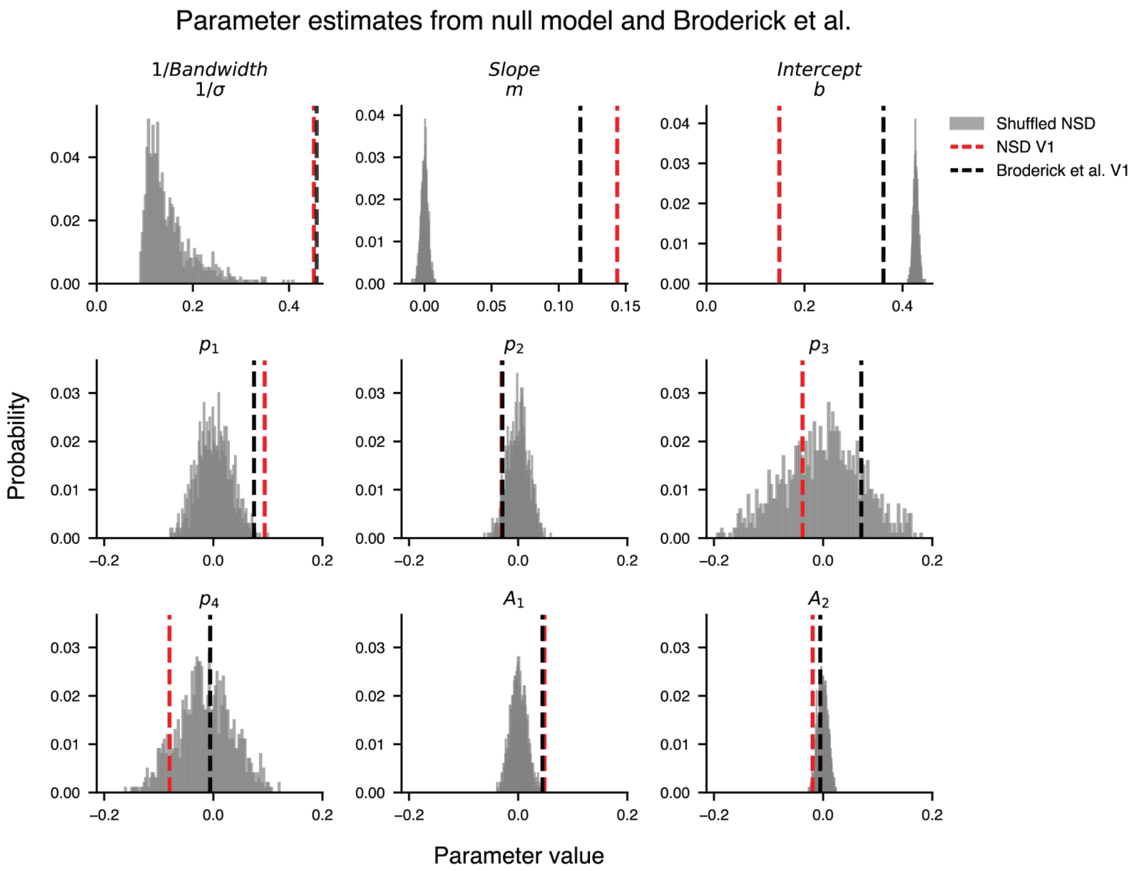


**Parameters from Broderick et al, NSD, and shuffled NSD**. Each panel is one of the nine parameters in the 2D spatial frequency model. The red and black lines are mean parameters estimates across participants from the NSD dataset and BRoderick et al, respectively. The gray bars represent histograms of the parameter estimates from shuffling the NSD data and refitting (see Methods, section 2.5. “Comparing models across datasets” for shuffling methods). The x-axis represents the parameter estimate without normalization.
